# Hybridisation has shaped a recent radiation of grass-feeding aphids

**DOI:** 10.1101/2022.09.27.509720

**Authors:** Thomas C. Mathers, Roland H. M. Wouters, Sam T. Mugford, Roberto Biello, Cock Van Oosterhout, Saskia A. Hogenhout

## Abstract

Aphids are common crop pests. These insects reproduce by facultative parthenogenesis involving several rounds of clonal reproduction interspersed with an occasional sexual cycle. Furthermore, clonal aphids give birth to live apterous or winged young that are already pregnant. Together, these qualities enable rapid population growth and have facilitated the colonisation of crops globally. In several cases so-called “super clones” have come to dominate agricultural systems. However, the extent to which the sexual stage of the aphid life cycle has shaped global pest populations has remained largely unclear, as have the origins of successful lineages. Here, we used chromosome-scale genome assemblies to disentangle the evolution of two of the most significant global pests of cereals – the English (*Sitobion avenae*) and Indian (*Sitobion miscanthi*) grain aphids. We found that genome-wide divergence between *S. avenae* and *S. miscanthi* is low and that *S. avenae sensu stricto* is part of a larger cryptic species complex that includes multiple diverged *S. miscanthi* lineages. Moreover, comparison of haplotype-resolved assemblies reveals that the *S. miscanthi* isolate used for genome sequencing is likely a hybrid, with one of its diploid genome copies being closely related to *S. avenae* (∼0.5% divergence) and the second being substantially more divergent (> 1%). Analyses of genotyping-by-sequencing (GBS) data of grain aphids from the UK and China revealed that *S. avenae* and *S. miscanthi* are part of a species complex with many highly differentiated lineages that predate the origins of agriculture. The complex consists of hybrid lineages that display a tangled history of hybridisation and genetic introgression. These data demonstrate that hybridisation has substantially contributed to grain aphid diversity, and hence, to the evolutionary potential of this important pest species.

## Introduction

Crop pests and pathogens have evolved from species colonising wild plants to take advantage of new niches created by agriculture (Stukenbrock and McDonald 2008; Bernal and Medina 2018). Often, these pests and pathogens can evolve rapidly to overcome pesticides, the introduction of resistant crop varieties or to subvert other control measures (McDonald and Stukenbrock 2016). Furthermore, pests and pathogens may periodically undergo host jumps causing disease outbreaks (Couch *et al*. 2005; Grünwald and Flier 2005; Inoue *et al*. 2017). The rapid evolution of pests and pathogens may occur via selection acting on standing genetic variation (Panini *et al*. 2021; Taylor *et al*. 2021; Pélissié *et al*. 2022), novel innovations derived from *de novo* mutation (Dong *et al*. 2014; Hartmann *et al*. 2021) or by acquiring genetic innovations from other species or populations through hybridisation and introgression (McMullan *et al*. 2015; Menardo *et al*. 2016; Valencia-Montoya *et al*. 2020; Rogério *et al*. 2022). Understanding the origins, diversity and evolutionary potential of pests and pathogens is therefore of fundamental importance.

Among insect crop pests, aphids – a diverse group of sap-sucking insects from order Hemiptera – are particularly important due to their role as vectors of plant disease agents (Ng and Falk 2006; Hogenhout *et al*. 2008; Whitfield *et al*. 2015). A key aspect of aphid success as crop pests is their “best of both worlds” approach to reproduction that involves multiple rounds of asexual reproduction alternated with occasional sexual reproduction (Dixon 1977; Moran 1992; Simon *et al*. 2010). This reproductive mode, known as cyclical parthenogenesis, involves specialised reproductive morphs and maximises population expansion via clonal propagation whilst providing opportunity for mixing of genetic variation during the sexual cycle. In spring and summer, asexual females rapidly produce live-born, genetically identical (with the exception of *de novo* mutation and gene conversion) offspring via apomictic parthenogenesis (Blackman 1979; Tomiuk and Wöhrmann 1982; Hales *et al*. 2002). Population expansions during this phase are further accelerated by telescoping of generations, whereby aphid females give birth to multiple live young (viviparity) that already have daughters developing inside them. Furthermore, winged asexually reproducing morphs may be generated that facilitate dispersal, potentially over long distances (Irwin *et al*. 1988). In the autumn and winter, differentiated male and female morphs are induced that reproduce sexually, producing overwintering eggs that hatch as asexual females the following spring, restarting the cycle. As such, unlike strictly sexual species, high fitness aphid genotypes fortuitously produced by sexual reproduction can be rapidly amplified during the asexual stage. In many cases this has led to the proliferation of so called “super clones” which dominate aphid populations and can spread globally (Haack *et al*. 2000; Figueroa *et al*. 2005; Peccoud *et al*. 2008; Margaritopoulos *et al*. 2009). However, the origins of highly successful aphid pest lineages are often unknown.

Cereal aphids present an ideal opportunity to investigate the evolutionary genomics of crop pest emergence, particularly given the global cultivation of wheat and other cereal crops. Specialisation on cereals and other grasses has occurred multiple times during aphid evolution and several grass-specialists have become important pests of crops (Blackman and Eastop 1984; Choe *et al*. 2006). Among aphid cereal-specialised lineages, the English and Indian grain aphids, *Sitobion avenae* and *Sitobion miscanthi*, are particularly destructive due to their global distribution and role as vectors of barley yellow dwarf virus (Vickerman and Wratten 1979). Interestingly, *S. avenae* and *S. miscanthi* have emerged from a species radiation. *Sitobion* is one of the most diverse genera of the subfamily Aphidinae, with at least 61 recorded species (Blackman and Eastop 1984; Favret 2022). Approximately half of these species colonise monocotyledons, with many specialising on grasses. Along with two other grain aphid species, *S. fragariae* and *S. akebiae*, the English and Indian grain aphids form a closely related complex (Choe *et al*. 2006). Across Europe and in China, population genetic studies have revealed high genetic diversity of *S. avenae* and *S. miscanthi* (Simon *et al*. 1999; Papura *et al*. 2003; Morales-Hojas *et al*. 2020). However, the genome-wide diversity and diversification of *S. avenae* and *S. miscanthi*, and their evolutionary origins, is currently unknown as studies have either focused on each lineage in isolation, or only made use of small number of microsatellite markers.

Here, we investigate the evolution of grain aphids from the *Sitobion* genus using chromosome-scale genome assemblies and population genomics. We generate chromosome-scale genome assemblies of *S. avenae* and a divergent grass-feeding aphid, *Rhopalosiphum padi* (bird cherry-oat aphid) and reassemble a recently published chromosome-scale genome sequence of *S. miscanthi* (Jiang *et al*. 2019). We also generated a high-quality draft genome assembly of *Metopolophium dirhodum* (rose-grain aphid), another important crop pest of grains, from a sister genus to *Sitobion* that serves as an outgroup in our analyses. On finding that *S. avenae* and *S. miscanthi* have low genome-wide sequence divergence and that the strain used to assemble the *S. miscanthi* genome is likely of hybrid origin, we reanalysed published population genomic data for *S. miscanthi* and *S. avenae* from the UK and China (Morales-Hojas *et al*. 2020). Using these data, we revealed that *S. avenae* and *S. miscanthi* are part of a larger cryptic species complex shaped by hybridisation.

## Results

### Chromosome-scale genome assemblies of Sitobion miscanthi and Sitobion avenae and a short-read assembly of Metopolophium dirhodum

We first assessed the recently published chromosome-scale genome assembly of *S. miscanthi* (Simis_v1) that derives from a Chinese lab colony dubbed Langfang-1 (Jiang *et al*. 2019). Simis_v1 was assembled with a combination of PacBio long-reads (85x coverage), Illumina short-reads (105x coverage) and *in vivo* Hi-C data (76x coverage) for long-range scaffolding (**Supplementary Table 1**). The total length of the assembly is 398 Mb, it has a contig N50 of 1.6 Mb and a scaffold N50 of 36.3 Mb, with the nine longest scaffolds in the assembly accounting for 95% of the assembled genome content (**Table 1**). These nine super-scaffolds are assumed to correspond to the nine chromosomes of *S. miscanthi* (Jiang *et al*. 2019).

**Table 1:** Genome assembly and annotation statistics.

| Species | *S. miscanthi* | *S. miscanthi* | *S. avenae* | *M. dirhodum* |
| --- | --- | --- | --- | --- |
| Assembly | Simis_v1 | Simis_v2 | Siave_v2.1 | Medir_v1.1 |
| Sequencing approach<sup>a</sup> | IL + PB + HiC | IL + PB + HiC | IL + HiC | IL |
| Base pairs (Mb) | 397.9 | 403.1 | 366.0 | 387.0 |
| % Ns | 0.01 | 0.13 | 0.80 | 0.08 |
| Number of contigs<sup>b</sup> | 1,148 | 1,889 | 23,024 | 28,240 |
| Contig N50 (Mb)<sup>b</sup> | 1.61 | 1.92 | 0.04 | 0.04 |
| Number of scaffolds | 655 | 833 | 14,626 | 24,973 |
| Scaffold N50 (Mb) | 32.70 | 37.52 | 29.84 | 0.05 |
| % of assembly in chromosome length scaffolds | 94.8 | 96.2 | 74.0 | NA |
| Protein coding genes | 16,006 | 21,798 | 19,919 | 22,349 |
| Transcripts | 16,006 | 23,875 | 22,368 | 24,826 |
| Reference | Jiang *et al.* 2019 | This study | This study | This study |

Alignment of Simis_v1 with chromosomes of the closely related model aphid *Acyrthosiphon pisum* (Mathers *et al*. 2021) reveals fragmentation of the X chromosome into three chunks as well as substantial rearrangement of the autosomes (**Supplementary Figure 1**). This is unexpected as we, and others, have recently shown long-term conservation of aphid X chromosome structure across divergent lineages (Li *et al*. 2020; Mathers *et al*. 2021). Fragmentation of the *S. miscanthi* X chromosome may represent genuine chromosome fission events or be the result of genome assembly error. To assess the quality of Simis_v1 chromosome-scale genomic scaffolds we generated a genome-wide Hi-C contact map using data from the original genome assembly (**Figure 1a**). Visual inspection of the Simis_v1 Hi-C contact map shows multiple off diagonal Hi-C contacts that are indicative of large-scale assembly error within scaffolds (Dudchenko *et al*. 2018; Howe *et al*. 2021). Furthermore, several scaffolds show regions that have very low contact frequencies with adjacent sequence in the scaffold, potentially indicating incorrect assignment of scaffold start and end points. Taken together, these analyses suggest that the assembled chromosomes of Simis_v1 are likely to be inaccurate, and that there may be only a single X chromosome in *S. miscanthi*.

**Figure 1:**
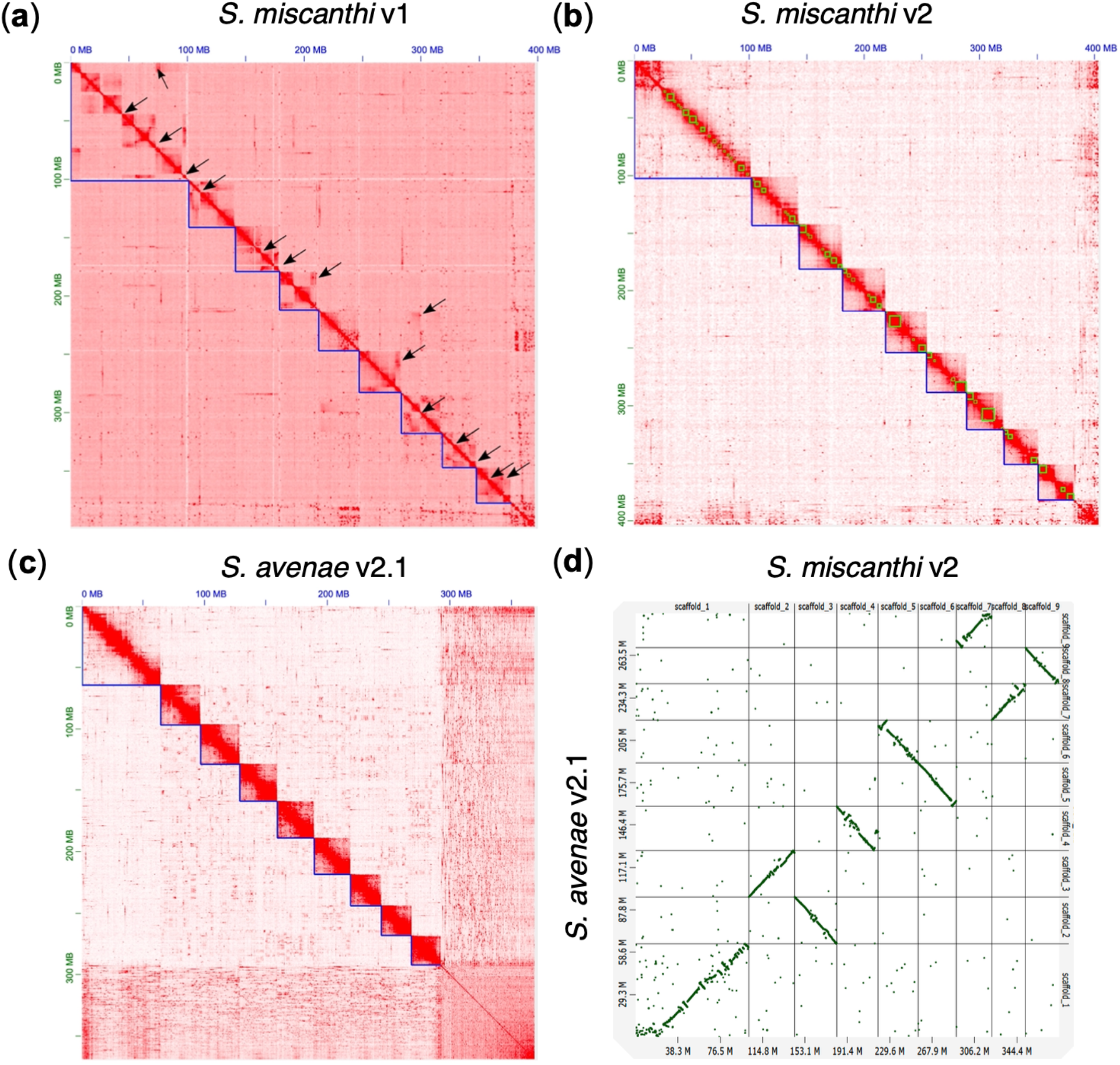
New chromosome-scale assemblies of *S. miscanthi* and *S. avenae* are high quality and show conserved synteny. (**a**) Hi-C contact map for the *S. miscanthi* v1 genome assembly. Blue lines show chromosome-scale super scaffolds. Arrows indicate likely assembly errors. Genomic scaffolds are ordered from longest to shortest with the x- and y-axis showing cumulative length in millions of base pairs (Mb). (**b**) Hi-C contact map for the *S. miscanthi* v2 genome assembly. Green lines show contigs. (**c**) Hi-C contact map for the *S. avenae* v2.1 genome assembly. (**d**) *Dot plot* showing a MashMap (Jain *et al*. 2018) whole genome alignment between the *S. miscanthi* v2 and *S. avenae* v2.1 genome assemblies. For clarity, only chromosome-scale scaffolds are included. Scaffolds in each assembly are ordered from longest to shortest. The x- and y-axis show cumulative scaffold length in Mb.

Given the apparent scaffolding errors in Simis_v1, we reassembled the *S. miscanthi* genome using the original sequence data to create Simis_v2. In total, Simis_v2 spans 403 Mb and 96% of the assembly is contained in nine chromosome-scale super scaffolds that have consistent Hi-C contact frequencies along their full length (**Table 1**; **Figure 1b**). Compared to Simis_v1, the contig N50 size of Simis_v2 is modestly improved (1.9 Mb vs 1.6 Mb; **Table 1**). Furthermore, based on the representation of arthropod Benchmarking sets of Universal Single-Copy Orthologs (BUSCOs; n=1,066), we substantially reduced the amount of missing (2.6% vs 5.3%) and duplicated (3.1% vs 4.7%) assembly content (**Supplementary Figure 2**). The improved sequence content of Simis_v2 compared to Simis_v1 is also supported by K-mer analysis of the raw Illumina reads with either genome assembly version (**Supplementary Figure 3**) and a taxon-annotated GC content-coverage plot indicates that Simis_v2 is free from obvious contamination (**Supplementary Figure 4**).

To further validate our new assembly of *S. miscanthi* and generate additional genomic resources for the *Sitobion* genus, we also generated a chromosome-scale assembly of the English grain aphid (*S. avenae*), using a combination of PCR-free Illumina short-read sequencing (75x coverage) and *in vivo* Hi-C (**Supplementary Table 1**). As we are currently assembling a diverse range of aphid species (Mathers *et al*. 2022), including several that are maintained at the John Innes Centre (JIC) Insectary, we experimented with using a mixed species sample to reduce Hi-C library preparation costs. We pooled *S. avenae* individuals with another aphid species – the bird cherry-oat aphid (*Rhopalosiphum padi*) – and sent the resulting pooled sample to Dovetail Genomics (Santa Cruz, CA) for Hi-C library preparation. As nuclei are cross-linked *in vivo* during Hi-C library preparation, species-specific chromatin conformation information is maintained for each species allowing a single library to be used to scaffold multiple species (Marbouty *et al*. 2014). Furthermore, high sequence divergence between *S. avenae* and *R. padi* (Aphidini vs Macrosyphini synonymous site divergence = ∼34% [Mathers et al. 2021]) minimises the chance of Hi-C reads mismapping between the two species. To support this effort, we also generated a new draft genome assembly from the *R. padi* clonal lineage maintained at JIC using 10x genomics linked reads.

In total, we generated 20.7 Gb of multi-species Hi-C data giving ∼10x coverage of the *S. avenae* genome and ∼30x coverage of the *R. padi* genome. The final assembly of *S. avenae* (Siave_v2.1) spans 369 Mb with a contig N50 size of 40 Kb (**Table 1**). BUSCO and K-mer analysis shows that the assembly is highly complete with little missing or duplicated content (**Supplementary Figure 2; Supplementary Figure 5**) and a taxon-annotated GC content-coverage plot indicates that Siave_v2.1 is free from obvious contamination (**Supplementary Figure 6**). The shorter total assembly size of Siave_v2.1 compared to Simis_v2 (369 Mb vs 403 Mb) is likely the result of missing and collapsed repeat content due to the use of short Illumina reads for *de novo* assembly. Despite high fragmentation, scaffolding with Hi-C enabled the placement of 74% of the *S. avenae* assembly onto nine chromosome-scale super scaffolds (**Figure 1c**). Whole genome alignment of Siave_v2.1 and Simis_v2 chromosome-scale scaffolds reveals broad structural agreement between the two independent assemblies, supporting the accuracy of our multi-species Hi-C scaffolding approach (**Figure 1d**). We also scaffolded our draft assembly of *R. padi* using the same multi-species Hi-C library, placing 95% of the draft assembly onto four chromosome-scale super scaffolds (**Supplementary Figure 7**) corresponding to the expected *R. padi* karyotype (2n = 8 [Monti *et al*. 2010]). We make our new *R. padi* assembly available here, but it will be described in more detail elsewhere.

Finally, to aid comparative genome analysis, we generated a short-read genome assembly of the rose-grain aphid, *Metopolophium dirhodum*, which is thought to be closely related to *Sitobion* (Choi *et al*. 2018). Following the low-cost genome assembly approach set out in Mathers *et al*. (2020) we generated 23.9 Gb (62x coverage) of PCR-free Illumina genome sequence data and 14.8 Gb of strand-specific RNA-seq data for genome assembly scaffolding and genome annotation (**Supplementary Table 1**). We assembled these data into 24,973 scaffolds spanning 387 Mb (Medir_v1.1; **Table 1**). Although fragmented (scaffold N50 = 49 Kb), this short-read assembly is highly complete at the gene level (BUSCO complete = 97.1%; **Supplementary Figure 2**) and K-mer analysis reveals the absence of excessive missing or duplicated genome content (**Supplementary Figure 8**). The assembly is also free from obvious contamination based on a taxon-annotated GC content-coverage plot (**Supplementary Figure 9**).

All three new grain aphid genome assemblies (Simis_v2, Siave_v2.1 and Medir_v1.1) were annotated using the same gene annotation pipeline that we previously applied to the model aphid species *A. pisum* and *Myzus persicae* (Mathers *et al*. 2021), incorporating evidence from RNA-seq data (**Supplementary Table 1**). In total, we annotated 21,798 genes (23,875 transcripts) in *S. miscanthi*, 19,919 genes (22,368 transcripts) in *S. avenae* and 22,349 genes (24,826 transcripts) in *M. dirhodum* (**Table 1**). We also annotated our new chromosome-scale assembly of *R. padi* using the same procedure, identifying 16,977 genes (19,137 transcripts). Compared to Simis_v1, our annotation of Simis_v2 identifies an additional 5,792 genes. This large increase in gene count is likely due to a combination of improved genome assembly completeness in Simis_v2 and the use of different gene annotation pipelines. Indeed, BUSCO analysis of the Simis_v1 and Simis_v2 gene sets reveals that Simis_v2 is substantially more complete than Simis_v1 (**Supplementary Figure 10**), with completely missing BUSCO genes reduced by 55% in Simis_v2 (n=68 vs n=28). Furthermore, RNA-seq pseudoalignment rates of the mixed whole-body sample of *S. miscanthi* from Jiang *et al*. (2019) to the annotated gene models increased from 78% in Simis_v1 to 83% in Simis_v2 (**Supplementary Table 2**). Taken together, our highly complete genome assemblies and annotations of *S. miscanthi, S. avenae* and *M. dirhodum* provide a solid foundation to study grain aphid biology and compliment two other contig-level long-read genome assemblies of *S. avenae* (clone SA3 [Byrne *et al*. 2022] and SaG1 [Villarroel *et al*. 2022]) that were published during the completion of this study.

### Genome evolution in dactynotine aphids

Increasing numbers of sequenced aphid genomes are allowing finer scale analysis of aphid genome evolution (e.g., Julca *et al*. 2019). To place our new assemblies of *S. miscanthi, S. avenae* and *M. dirhodum* in a phylogenetic context we clustered their proteomes with eight other aphid species from the aphid tribes Macrosiphini and Aphidini (**Supplementary Table 3**). In total, we clustered 270,894 proteins into 23,712 orthogroups (gene families) and 19,653 singleton genes (**Supplementary Table 4**). Maximum likelihood phylogenetic analysis based on a concatenated alignment of 5,091 conserved single-copy genes produced a fully resolved species tree that places *Sitobion* and *Metopolophium* in a monophyletic group that is closely related to *A. pisum* (**Figure 2a**; **Supplementary Figure 11**). These findings are in agreement with previous studies using small numbers of loci and larger sets of taxa (von Dohlen *et al*. 2006; Choi *et al*. 2018).

**Figure 2:**
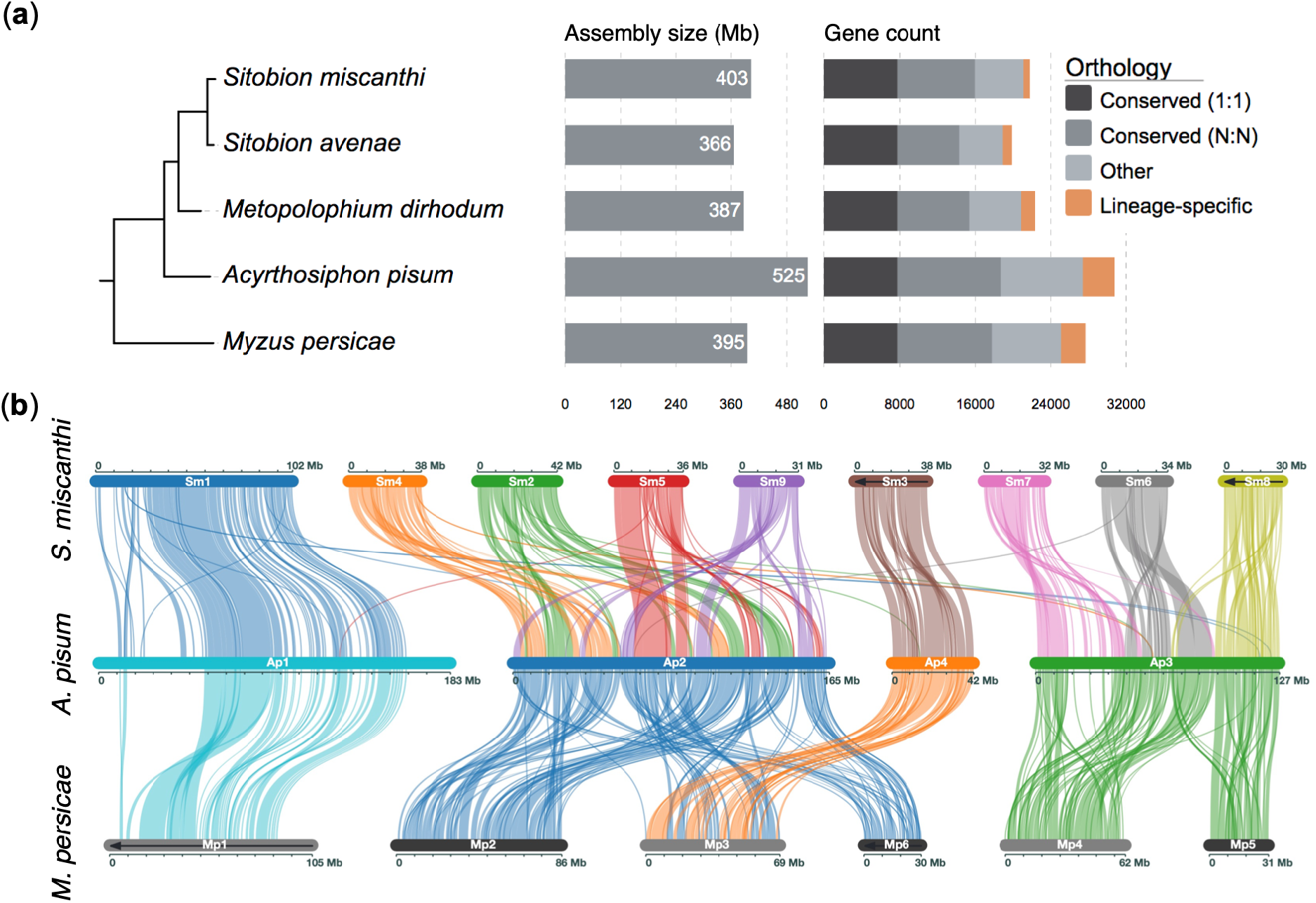
Comparative genomics of the dactynotine sub-tribe. (**a**) Maximum likelihood phylogeny of dactynotine aphids based on a concatenated alignment of 5,091 conserved single-copy protein coding genes. We included seven outgroup aphid species from Aphidinae but for simplicity only show *M. persicae*. The full phylogeny is shown in **Supplementary Figure 11**. All branches received maximal support according to the Shimodaira-Hasegawa test (Shimodaira and Hasegawa 1999) implemented in FastTree (Price *et al*. 2009, 2010) with 1,000 resamples. Branch lengths are in amino acid substitutions per site. The tree is annotated with genome assembly length and gene counts coloured by orthology relationships across the full phylogeny. (**b**) Chromosome evolution in the dactynotine sub-tribe. Plot shows blocks of syntenic genes identified between *S. miscanthi* (top), *A. pisum* (middle) and *M. persicae* (bottom) chromosomes. Chromosomes containing black arrows are visualised as the reverse compliment to aid clarity. Ap1 and Mp1 in *A. pisum* and *M. persicae*, respectively, have previously been identified as the X (sex) chromosome and are homologous to Sm1 (scaffold_1) in our new *S. miscanthi* (Simis_v2) genome assembly.

Assembly of three genomes closely related to the model aphid *A. pisum* – all belonging to the dactynotine sub-tribe (Börner and Heinze 1957; von Dohlen *et al*. 2006) – allows additional insights into gene family dynamics in aphids. *A. pisum* was the first aphid species to have its genome sequenced and this effort revealed substantial gene family expansion and, at the time, the largest gene count of any sequenced insect species (IAGC 2010). Subsequent reassembly of *A. pisum* has led to refinement of the estimated gene count for this species, supporting the large number of genes (Li *et al*. 2019; Mathers *et al*. 2021). Here, we find lower gene counts (19,919 – 22,349 vs 30,784) and smaller genome sizes (366 – 403 Mb vs 525 Mb) in *Sitobion* and *Metopolophium* compared to *A. pisum*, indicating that genome expansion is limited to the *A. pisum* lineage and occurred after divergence of the dactynotine common ancestor (**Figure 2a**).

In addition to dynamic gene family evolution, we and others have identified high rates of autosomal chromosome rearrangement in aphids (Li *et al*. 2020; Mathers *et al*. 2021). To investigate chromosome evolution in *Sitobion* and their dactynotine relatives, we identified syntenic genome regions between Simis_v2 and the chromosome-scale assemblies of *A. pisum* and *M. persicae* using MCscanX (Wang *et al*. 2012). This analysis confirms homology (and conservation) of the aphid X chromosome within Macrosiphini and reveals substantial autosomal genome reorganisation over the course of dactynotine aphid diversification (**Figure 2b**).

So far, the high rate of autosome rearrangement and small number of chromosome-scale genome assemblies have hampered the inference of specific chromosome rearrangement events that have led to extant aphid karyotypes (Mathers *et al*. 2021). However, we previously hypothesised that *A. pisum* chromosome 3 (Ap3) was formed by a fusion event involving homologs of *M. persicae* chromosomes 4 (Mp4) and 5 (Mp5) (Mathers *et al*. 2021). This scenario is confirmed by alignment of our new *S. miscanthi* assembly with *A. pisum* and *M. persicae*, with *S. miscanthi* chromosome 8 (Sm8) and *M. persicae* chromosome 5 (Mp5) sharing synteny with the final third (orientation as per the *A. pisum* JIC1 assembly) of Ap3 (**Figure 2b**). Additionally, the alignment of Mp4 and the first two thirds of Ap3 to *S. miscanthi* chromosome 7 (Sm7) and chromosome 6 (Sm6) reveals that Sm7 and Sm6 were formed by a chromosome fission event in the *Sitobion* lineage. As such, we can infer at least one unambiguous chromosome fusion event in the *A. pisum* lineage and one unambiguous chromosome fission event in the *Sitobion* lineage. Interestingly, we also find that the two large autosomes found in pea aphid (Ap2 and Ap3) exhibit distinct rearrangement patterns. Ap2 is homologous to four small (42 – 31 Mb) chromosomes in *S. miscanthi*, the order of which is shuffled in *A. pisum*. In contrast, Ap3 is homologous to Sm6, Sm7 and Sm8, which each align to distinct territories along Ap3. In the future, additional chromosome-scale assemblies of *S. miscanthi* and *A. pisum* close relatives will further illuminate the complex history of aphid chromosome evolution.

### Low genome-wide divergence between S. miscanthi and S. avenae

Phylogenetic analysis revealed a short branch length between *S. miscanthi* and *S. avenae* indicating recent divergence from a common ancestor (**Figure 2a**). To further investigate genome-wide patterns of sequence divergence among *Sitobion* aphids and their relatives we generated a reference-free, four-way, whole genome alignment between *S. miscanthi, S. avenae, M. dirhodum* and *A. pisum* using Progressive Cactus (Armstrong *et al*. 2020). Alignment coverage of our highly complete long-read-based *S. miscanthi* genome assembly ranged from 72% to 93% (**Figure 3a panel 1**) whereas alignment coverage of *S. avenae* was slightly higher (75% to 97%; **Figure 3a panel 2**), likely due to lower representation of hard to align repetitive regions in this short-read assembly. Using these alignments, we estimated pairwise sequence divergence in 1 Mb fixed windows along *S. miscanthi* chromosome-scale scaffolds (**Figure 3b**). We used alignments anchored to autosomal chromosomes to estimate genome-wide sequence divergence as aphid X chromosomes are known to exhibit elevated substitution rates (Jaquiéry *et al*. 2012, 2018). Using these autosomal alignments, we estimate median sequence divergence between *S. miscanthi* and *S. avenae* to be only 0.78% (**Figure 3c**). In contrast, *S. miscanthi* divergence from *M. dirhodum* and *A. pisum* is 5.63% and 8.86%, respectively.

**Figure 3:**
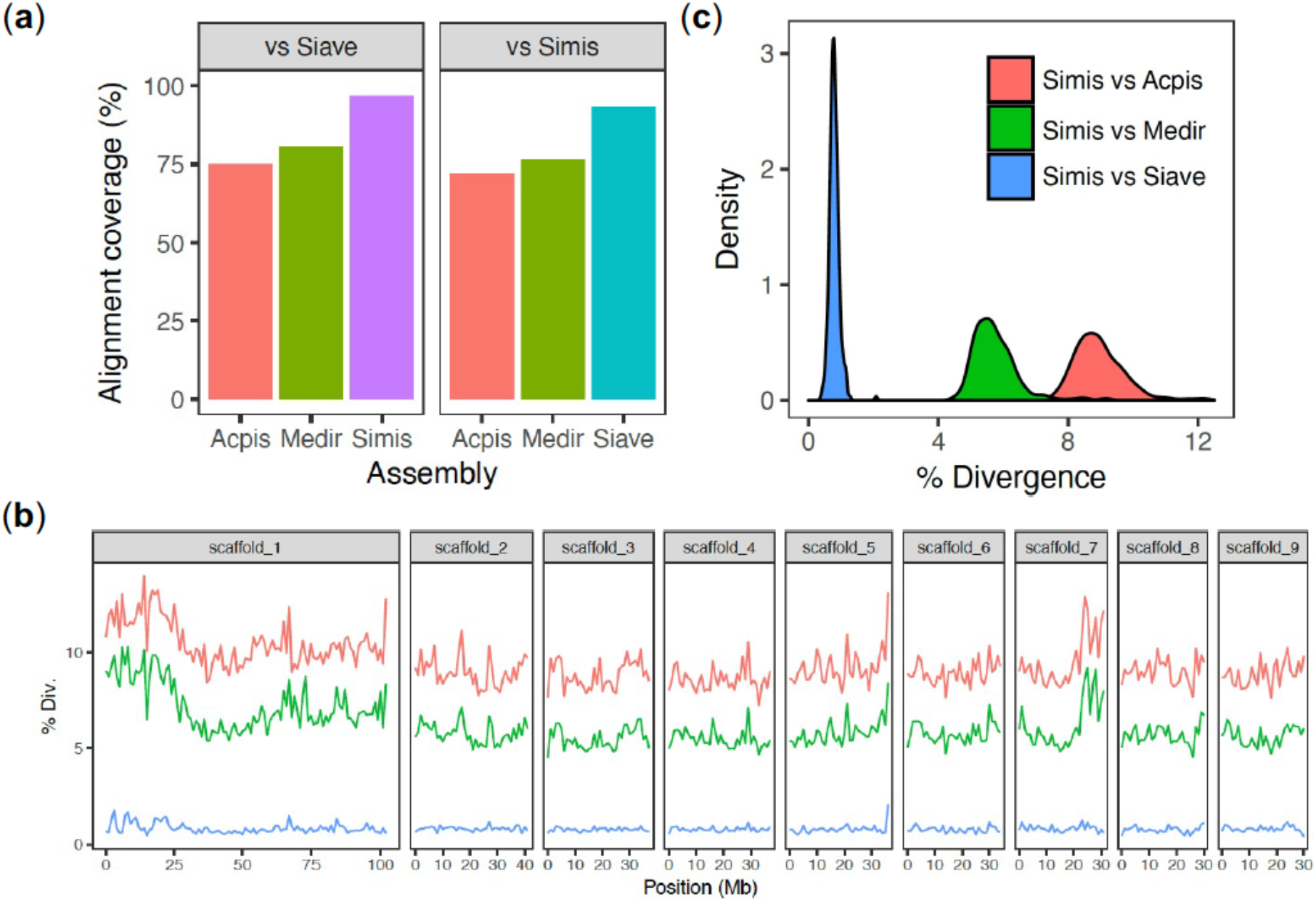
Reference-free whole genome alignment reveals low sequence divergence between *S. miscanthi* and *S. avenae*. (**a**) Alignment coverage on either *S. avenae* (Siave) or *S. miscanthi* (Simis) of the *A. pisum* (Acpis), *M. dirhodum* (Medir) and either *S. miscanthi* (Simis) or *S. avenae* (Siave) genome assemblies. (**b**) Pairwise sequence divergence between *S. miscanthi* and either *A. pisum* (red line), *M. dirhodum* (green line) or *S. avenae* (blue line) in 1 Mb fixed windows along the *S. miscanthi* genome assembly. scaffold_1 = the X (sex) chromosome. (**c**) *Density plot* showing the distribution of pairwise sequence divergence (as in **b**) for *S. miscanthi* autosomes vs those of *A. pisum* (median divergence = 8.86%), *M. dirhodum* (median divergence = 5.63%) and *S. avenae* (median divergence = 0.78%).

### Phased genome assemblies reveal hybrid origins of the S. miscanthi Langfang-1 lab population

Intriguingly, we noticed that the genome-wide divergence between *S. miscanthi* and *S. avenae* (∼0.78%) is lower than the predicted diversity (i.e. heterozygosity) found within the previously sequenced *S. miscanthi* clonal lineage dubbed Langfang-1 (LF1) (0.98% based on k-mer analysis of short-reads [Jiang *et al*. 2019]). To further investigate intra- and inter-individual patterns of sequence divergence in *S. miscanthi* and *S. avenae* we reconstructed independent phased haplotypes for each assembly and generated a four-way whole genome alignment with sibeliaZ (Minkin and Medvedev 2020). In total, we phased 3,043,224 (>99.99%) and 1,033,184 (97.68%) heterozygous single nucleotide polymorphisms (SNPs) and small indel variants on *S. miscanthi* (LF1) and *S. avenae* (JIC1) chromosomes, respectively. Due to the inclusion of long-range phase information from our Hi-C data, a single-phase block covered >=99.96% of each chromosome in both JIC1 and LF1 (**Supplementary Table 5**).

Using our whole genome haplotype alignment, we estimated pairwise divergence in 100 Kb fixed windows along the *S. avenae* reference genome. Genome-wide within-individual haplotype divergence (heterozygosity) is significantly lower in JIC1 than in LF1 (0.32% ± 0.0039% [mean ± *SE*] vs 0.83% ± 0.016%; Wilcoxon signed-rank test, *p* < 2.2 × 10^−16^; **Figure 4a**). This is possibly caused by an extreme founder event and/or inbreeding in the UK *S. avenae* population (Morales-Hojas *et al*. 2020). Consistent with this, we find mega base-scale stretches of near-zero haplotype divergence (i.e., long runs of homozygosity) on several *S. avenae* JIC1 chromosomes (**Figure 4b**; **Supplementary Figure 12**). Surprisingly, the two haplotypes found within LF1 substantially differ in their divergence from both JIC1 haplotypes, with one haplotype diverged by ∼0.5% and the other by ∼1.0% (**Figure 4a**). This unusual pattern of haplotype divergence is maintained across most chromosomes without any “switching” of haplotypes that would be expected if recombination had taken place (**Figure 4b; Supplementary Figure 13)**. As such, we hypothesise that the LF1 clonal lineage is a “frozen hybrid”, in particular, a first-generation (F1) clonal descendant of a cross between two lineages that differ in their divergence from *S. avenae*.

**Figure 4:**
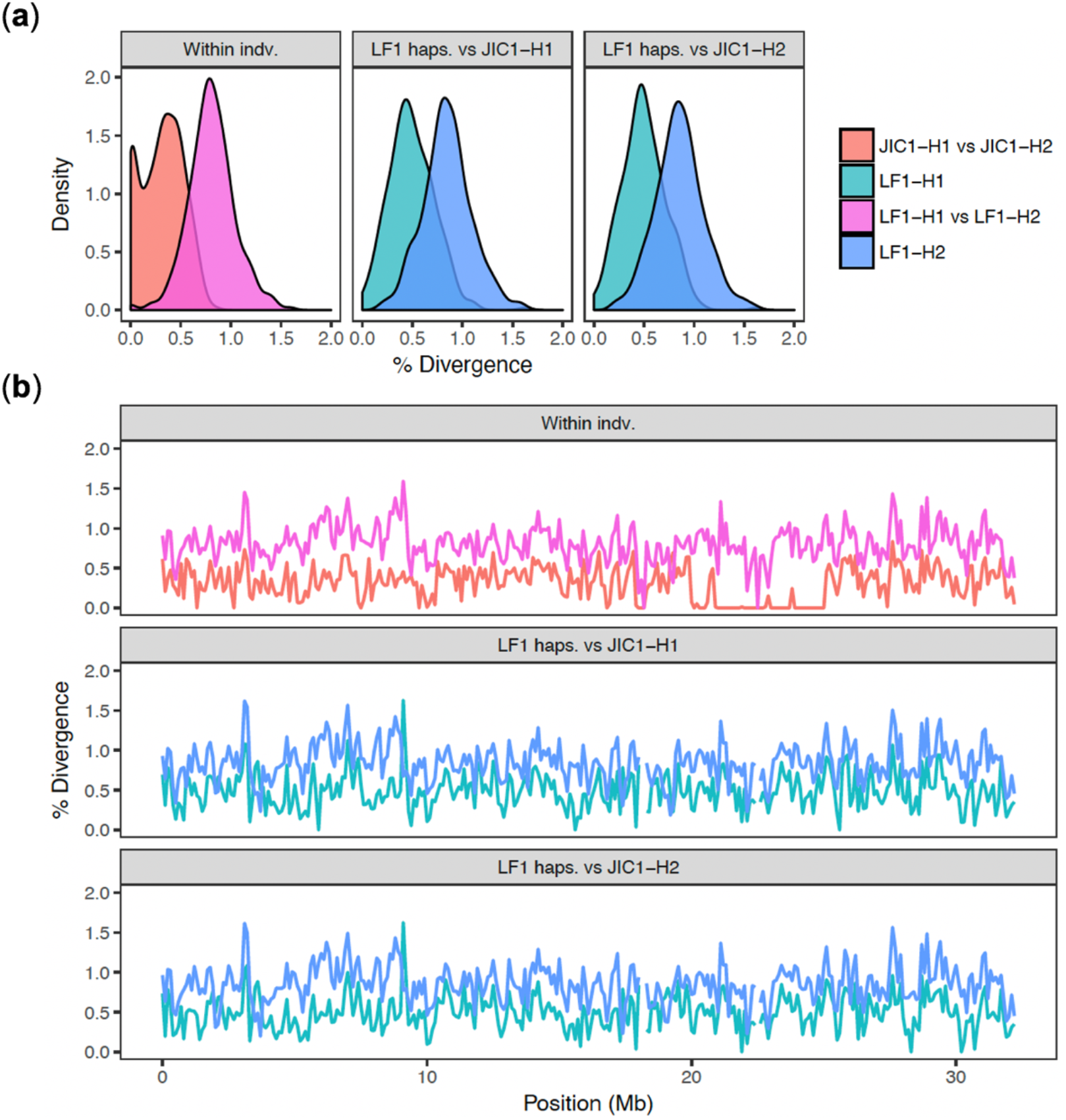
Highly differentiated haplotypes within the *S. miscanthi* LF1 clonal lineage differ in their in their divergence from *S. avenae*. (**a**) Sequence divergence distribution within and between phased haplotypes of JIC1 and LF1 for *S. avenae* chromosome (scaffold) 2. Comparing the LF1 haplotypes (LF1-H1 and LF1-H2) to either JIC1 haplotype reveals distinct patterns of divergence, with LF1-H1 diverged ∼0.5% from either JIC1 haplotype, and LF1-H2 diverged by ∼1%. (**b**) Haplotype divergence within and between JIC1 and LF1 in 100 Kb fixed windows along *S. avenae* chromosome (scaffold) 2. The unusual pattern of haplotype divergence observed in LF1 is maintained across the full length of chromosome 2 indicating an absence of recombination. Divergence patterns for all chromosomes are shown in **Supplementary Figure 12**.

### S. miscanthi and S. avenae are part of a cryptic species complex

To investigate the origins of the LF1 clone and gain a greater understanding of population-level divergence and diversity in *S. avenae* and *S. miscanthi* we reanalysed genotyping by sequencing (GBS) data for 100 *S. miscanthi* individuals from China and 119 *S. avenae* individuals from the UK from a recent study by Morales-Hojas *et. al*. (2020) (**Supplementary Table 6**). Previously, these data were analysed separately using different GBS protocols. However, given our finding that sequence divergence between *S. avenae* and *S. miscanthi* is very low, we reanalysed these data together and searched for overlapping SNP markers between the two sets of samples. In total, including variants from the JIC1 and LF1 whole genome samples, we identified 3,246,566 biallelic SNPs (min. depth >= 2 and site quality >= 30). Of these sites, we retained 4,359 that were covered (and called) in at least 75% of samples. We further refined the dataset by removing 60 samples that had more that 30% missing data. The final dataset contains markers spread across all nine *Sitobion* chromosomes (n = 332-978 per chromosome; **Supplementary Table 7**) and includes 149 samples, 52 from the UK and 97 from China, allowing us to investigate diversity and differentiation within and between *S. avenae* and *S. miscanthi* populations.

Previously, Morales-Hojas *et. al*. (2020) identified six highly differentiated *S. miscanthi* populations in China (genome-wide FST = 0.13 – 0.79). These results are recapitulated in our analysis using the shared SNP set, with highly similar groupings based largely on geographic location (**Figure 5a**). Furthermore, the LF1 and JIC1 whole genome samples cluster within their expected populations based on geography, i.e., LF1 groups with the Langfang Chinese *S. miscanthi* samples and JIC1 groups with the UK *S. avenae* samples. Surprisingly, by combing data for *S. avenae* and *S. miscanthi*, we find that the UK *S. avenae* population is more closely related to one of the Chinese *S. miscanthi* populations (TG_YC). This suggests that *S. avenae sensu stricto* may be part of a larger cryptic species complex that includes multiple diverged *S. miscanthi* lineages.

**Figure 5:**
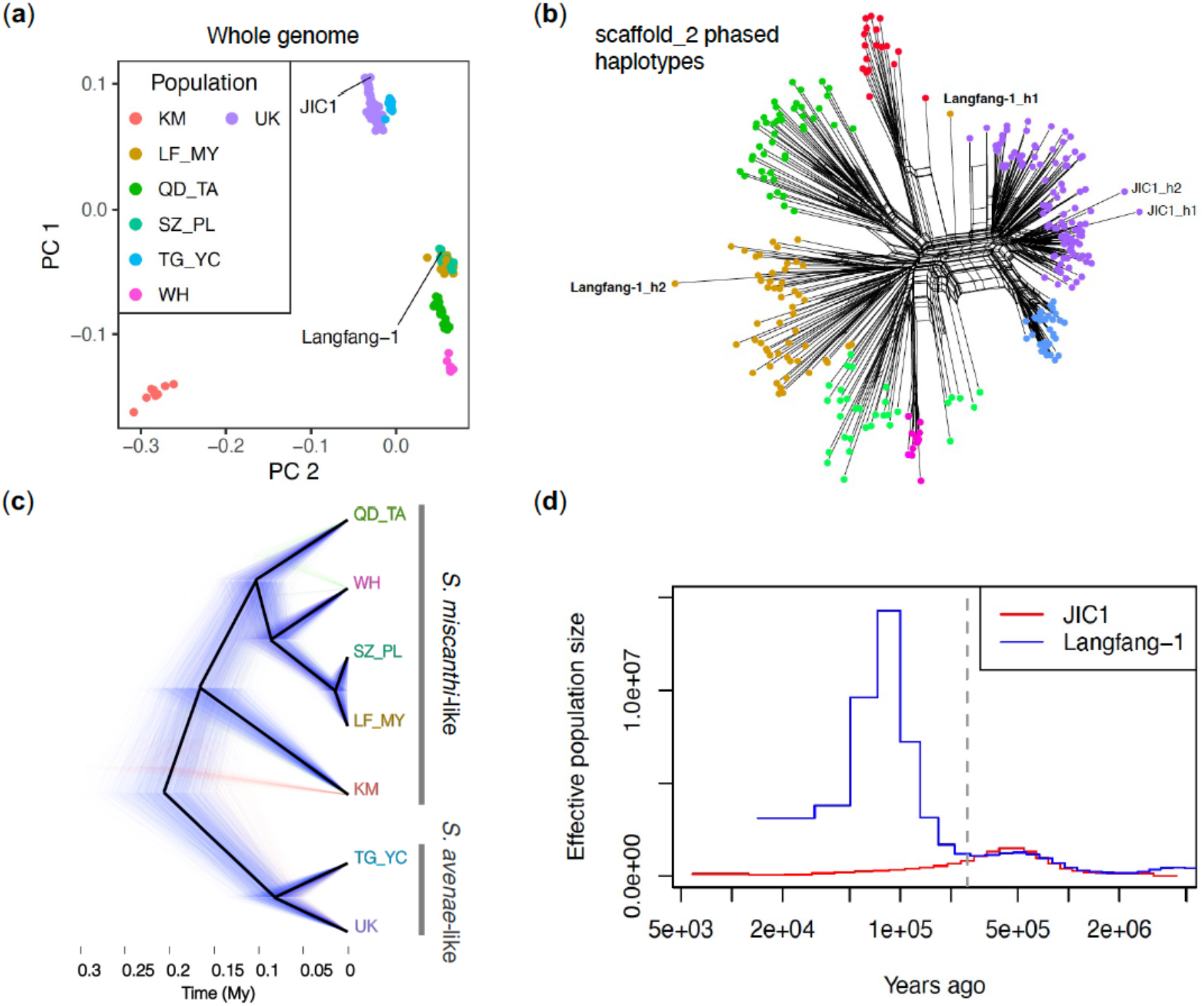
Population genomics of *Sitobion avenae* and *Sitobion miscanthi* from the UK and China. (**a**) Principle component analysis based on the thinned (max 1 SNP per 25Kb) shared SNP set (n = 1,772) reveals previously described population structure in China (Morales-Hojas *et al*. 2020) and the close relationship of the UK S. *avenae* (purple dots) to a Chinese *S. miscanthi* lineage (TG_YC; light blue dots). Samples used for genome assembly of *S. avenae* (JIC1) and *S. miscanthi* (LF1) are highlighted. Populations are named following Morales-Hojas *et al*. (2020) according to geographic location: KM = Kunming, LF_MY = Langfang and Mianyang, QD_TA = Qingdao and Tai’an, SZ_PL = Suzhou and Pingliang, TG_YC = Taigu and Yinchuan, WH = Wuhan, UK = United Kingdom. (**b**) SplitsTree network of samples shown in (**a**) based on phased haplotypes for SNPs on *S. aveane* chromosome 2. The two haplotypes from the *S. avenae* (JIC1) and *S. miscanthi* (Langfang-1) samples used for genome assembly are highlighted. (**c**) “Cloudogram” showing a time calibrated phylogeny of *Sitobion* lineages rooted with *M. dirhodum* (not shown). A posterior sample of 1,801 trees are drawn with the first, second and third most common topologies coloured blue, green, and red, respectively. Black lines show the maximum-clade-credibility tree. Tip labels (showing populations) are coloured according to (**a**). (**d**) Demographic history of the JIC1 (derived from the UK population) and LF1 (derived from the LF_MY population) whole genome sequence isolates estimated with MSMC2. The dashed vertical line indicates the approximate time of divergence between the two samples 250,000 years ago (Kya) and corresponds to the deepest split shown in (**c**) between the *S. avenae*-like and *S. miscanthi*-like lineages. Before 250 Kya, JIC1 and Langfang-1 have a shared demographic history.

To gain more insight into the putative hybrid origin of the *S. miscanthi* LF1 sample used for *de novo* genome assembly (**Figure 4**), we phased the shared SNP set using beagle (Browning and Browning 2007) and estimated a SplitsTree (Huson and Bryant 2006) network of the resulting haplotypes using data from *S. avenae* chromosome 2 (**Figure 5b**). This analysis reveals that one LF1 haplotype clusters with the Langfang *S. miscanthi* population and that the second falls out on its own between the monophyletic groups containing the UK *S. avenae*-like samples and the *S. miscanthi* KM population. As such, the LF1 clone is likely the product of a hybridisation event between the Langfang population and another, as yet unsampled, *Sitobion* lineage.

Next, we set out to date the radiation of the *Sitobion avenae* / *miscanthi* complex. The spontaneous mutation rate of the closely related aphid *A. pisum* has recently been inferred, enabling the estimation of species and population divergence times from genome sequence data (Fazalova *et al*. 2020). However, our (primarily) GBS-based SNP dataset does not include invariant sites and so likely suffers from ascertainment bias, making direct dating based on a known mutation rate challenging (Stange *et al*. 2018). To avoid applying an unrealistic mutation rate to our SNP data, we first estimated the divergence time of the sister genera *Sitobion* and *Metopolophium* to provide a calibration point for estimating divergence times within *Sitobion*. Using 8,462 autosomal phylogenetically inferred single-copy orthologs between *S. miscanthi* and our new genome sequence of the rose-grain aphid (*M. dirhodum*), we estimated the most recent common ancestor (MRCA) of *Sitobion* and *Metopolophium* to have accrued 3.64 million years ago (Mya) based on median autosomal synonymous site divergence of 5.9% (**Supplementary Figure 14**) and the *A. pisum* spontaneous mutation rate of 2.7 × 10^−10^ per haploid genome per generation and an average of 15 generations (14 asexual and 1 sexual) per year (Loxdale and Balog 2018). We then mapped genomic reads for *M. dirhodum* to the *S. avenae* reference genome and called SNPs alongside the *Sitobion* GBS and whole genome sequence samples, identifying 3,043 shared biallelic SNPs.

Using the SNP dataset and calibration point inferred above, we jointly estimated a population tree and divergence times under the multispecies coalescent (MSC) with SNAPP (Bryant *et al*. 2012) following recommendations by Stange *et al*. (2018). This analysis recovers a well-supported tree (**Figure 5b**; **Supplementary Figure 15)** that places the UK *S. avenae*-like population in a monophyletic group with the Chinese TG_YC population (Bayesian posterior probability (BPP) = 1) and the remaining *S. miscanthi*-like Chinese populations in second monophyletic group (BPP = 0.99). The deep split between the *S. miscanthi*-like and *S. avenae*-like groups is inferred to have occurred around 199 thousand years ago (Kya) (95% highest posterior density (HPD) = 141 – 263 Kya). Within the *S. miscanthi*-like group, the KM population forms a third highly differentiated lineage that diverged around 159 Kya (95% HPD = 108 – 206 Kya), although there is some support for an alternative topology which places this group as an outgroup to all other included *Sitobion* lineages pushing back the inferred split time. Within the *S. avenae*-like lineage, we estimate that the UK lineage diverged from the Chinese TG_YC lineage around 81 Kya (95% HPD = 51 – 111 Kya).

To provide an independent estimate of the primary split time between the *S. avenae*-like lineage and the *S. miscanthi*-like lineage based on whole genome sequences (rather than GBS samples), we estimated the past demographic history of the JIC1 and LF1 clonal lineages using MSMC2 (Malaspinas *et al*. 2016) which implements the multiple sequentially Markovian coalescent (MSMC) model. This analysis indicates that JIC1 and LF1 have a shared demographic history until around 250 Kya (**Figure 5d**), consistent with the divergence time between the *S. avenae*-like and *S. miscanthi*-like lineages estimated by SNAPP (**Figure 5c**). Furthermore, the LF1 sample shows a very sharp rise in effective population size around 100 Kya in (**Figure 5d**). Typically, in MSMC models of hybrids, the effective population size goes to infinity at the time when gene flow stopped between the parental lineages (Li and Durbin 2011; Cahill *et al*. 2016). The sharp rise in effective population size may therefore represent the divergence time of the LF_MY lineage from the second unidentified lineage that contributed to the LF1 hybrid. The failure to reach infinity may reflect a complex history of admixture between the parental lineages, or backcrossing following the initial hybridisation event. This could retain a signal of coalescence in small parts of the genome, stopping MSMC from estimating an infinite effective population size. Additional sequencing of Chinese *Sitobion* populations will likely shed further light on these processes. Nonetheless, our analyses reveal multiple highly differentiated lineages within the *S. miscanthi* / *avenae* complex that substantially predate the origins of agriculture.

### Hybridisation has shaped the Sitobion radiation

Finally, given the putative hybrid origins of the *S. miscanthi* LF1 lab population we asked whether hybridisation has occurred more widely between lineages in the *S. avenae* / *miscanthi* complex. Using *M. dirhodum* as an outgroup, the SNAPP species/population tree, and excluding the previously identified Langfang-1 hybrid isolate, we summarised admixture across the *S. avenae* / *miscanthi* complex using Patterson’s *D* (ABBA–BABA test) (Patterson *et al*. 2012) and the *f*-branch (*f*_b_) statistic (Malinsky *et al*. 2018), both of which use patterns of allele sharing to infer gene flow. First, to gain a course and conservative overview of admixture history across the complex, we calculated the minimum *D* statistic (*D*_min_) irrespective of phylogeny for all trios of ingroup lineages (n=35) (Malinsky *et al*. 2018). In total, 34% of trios have significant *D*_min_ (Bonferroni corrected *p* < 0.05; **Supplementary Table 8**), indicating moderate levels of admixture among members of the *S. avenae* / *miscanthi* complex.

Significant *D*_min_ may be caused by current admixture (i.e., hybridisation) between extant taxa, or by historical admixture between ancestral lineages. Historic admixture is expected to inflate the number of trios with significant *D*_min_ due to non-independence. To map admixture to specific lineages of the *S. avenae* / *miscanthi* complex we calculated the *f*_b_ statistic which summarises all possible *f*_4_ admixture ratios for a given phylogeny and reports the admixture proportion between all compatible pairs of branches and taxa in a phylogeny (Malinsky *et al*. 2018). We find the largest admixture proportions between WH and the QD_TA (*f*_b_ = 0.57) and KM (*f*_b_ = 0.28) lineages (**Figure 6; Supplementary Table 8**). There are also strong bidirectional signatures of admixture between the *S. avenae*-like group and the *S. miscanthi*-like group. In particular, admixture is detected between the TG_YC *S. avenae*-like lineage in China and among all members of the *S. miscanthi*-like group (*f*_b_ = 0.07 – 0.29; LF_MY strongest), and also between the common ancestor of *S. avenae*-like LF_MY + SZ_PL. This latter signal may reflect admixture event(s) prior to the divergence of the two sampled *S. avenae*-like lineages. However, better sampling and an improved understanding of global *S. avenae / S. miscanthi* diversity and phylogeography will be required to better understand specific gene flow events within the complex.

**Figure 6:**
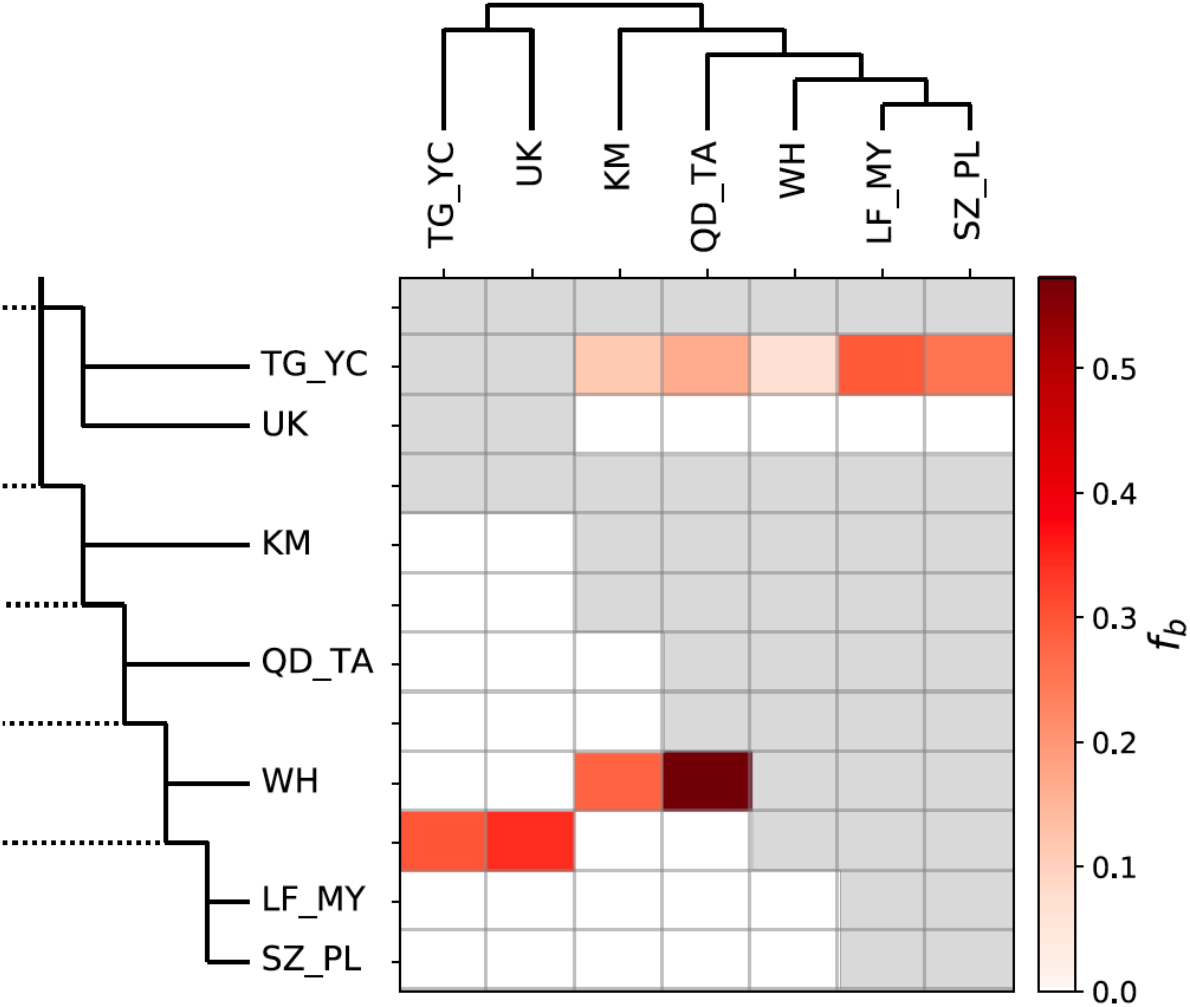
Excess allele sharing between *Sitobion* lineages. The *heatmap* shows the magnitude of the *f*_b_ ratio (Malinsky *et al*. 2018) between each branch on the y-axis and the sample on the x-axis. Grey squares indicate comparisons that cannot be made. Comparisons where corresponding *D* statistics (**Supplementary Table 8**) are non-significant (*p* > 0.01) are set to zero. Tip names correspond to populations/lineages identified in **Figure 5a**.

Next, to investigate admixture across the *S. avenae* / *miscanthi* complex at greater resolution, we generated a second SNP dataset containing only the Chinese GBS samples and the two whole genome sequences of Langfang-1 and JIC1 (n=98). As all the Chinese samples derive from the same GBS experiment, we were able to recover a much larger set of SNPs enabling fine-scale introgression analysis across the genome. In total, we identified 73,903 SNPs that are present in at least 90% of samples. A phylogenetic network based on this larger SNP set reveals large reticulations indicating substantial admixture (or hybridisation) between lineages of the *Sitobion miscanthi* / *avenae* complex (**Figure 7a**), consistent with our previous analysis based on the smaller SNP set (**Figure 6**).

**Figure 7:**
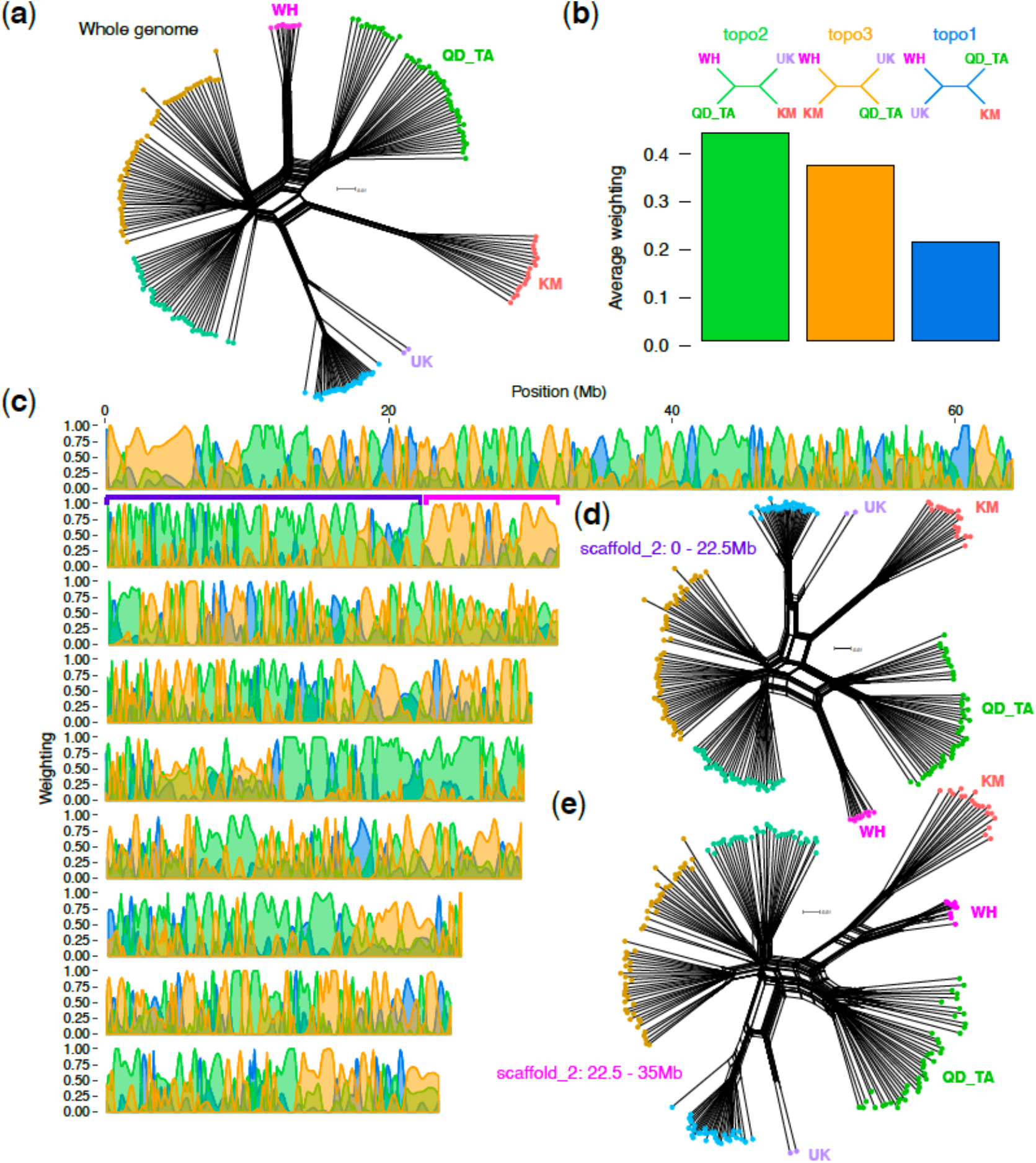
Topology weighting reveals massive introgressed blocks. (**a**) SplitsTree (Huson and Bryant 2006) network of phased haplotypes based on 73,903 genome-wide SNPs for Chinese GBS samples (**Supplementary Table 6**) plus the whole genome sequences of *S. avenae* JIC1 from the UK and *S. miscanthi* LF1 from the LF_MY population. Tips are coloured by population/lineage as per **Figure 5**. Three focal *S. miscanthi*-like lineages (WH, QD_TA and KM) and one outgroup *S. avenae* lineage (UK) are highlighted. (**b**) Phylogenetic trees were estimated in 50 SNP windows across all *S. avenae* chromosomes for all samples included in (**a**). The *bar chart* summarises the average genome-wide weighting (determined by Twisst [Martin and Van Belleghem 2017], see *main text*) of the three possible topologies between WH, KM, QD_TA and UK focal lineages. The tree with the highest weighting (topo2) corresponds to the species tree estimated with SNAPP (**Figure 5c**). Alternative topologies are also highly weighted. (**c**) The distribution of topology weightings across all nine *S. avenae* chromosomes reveals large blocks with a different evolutionary history to the species tree. Example regions on chromosome (scaffold) 2 are highlighted with purple (predominantly weighted towards the species tree (topo2)) and pink (predominantly weighted towards topo3). (**d**) and (**e**) show SplitsTree networks of phased haplotypes for the example regions of chromosome 2 highlighted in (**c**). For each network, the focal lineages are indicated.

Using the large SNP set we investigated genome-wide patterns of introgression focussing on the WH, QD_TA and KM lineages, which are estimated to have the highest rates of hybridization based on the *f*_b_ statistic (**Figure 6**). We estimated phylogenetic trees in 50 SNP windows for all samples and summarised the distribution of all possible topologies among the three focal lineages and the UK *S. avenae* lineage using topology weighting by iterative sampling of subtrees (Twisst: Martin and Van Belleghem 2017). As expected, Twisst recovers the SNAPP species topology as the highest weighted topology genome-wide (topo2: **Figure 7b**) – this tree places the KM lineage as a sister to the WH and QD_TA lineages. However, the second most common topology (topo3) also receives high weighting and groups the KM and WH lineages together with QD_TA as an outgroup (**Figure 7b**). This pattern is consistent with admixture between the WH and KM lineages (or their ancestors).

Surprisingly, the distribution of the three topologies across the genome is non-random with large regions of each chromosome predominantly having either the species tree (topo2) or the admixture tree (topo3) (**Figure 6c**). A striking example is found on chromosome 2 where a clear switch point can be seen at ∼22.5Mb, with the beginning of the chromosome (*region 1*) strongly weighted towards the species tree, and the final 12.5 Mb (*region 2*) strongly weighted towards the admixture tree. Network analysis based on SNPs from either of these two regions shows that the KM individuals group closely to the *S. avenae*-like group (UK + TG_YC) in *region 1* (**Figure 7d**), but that they group more closely to the WH individuals in *region 2* (**Figure 7e)**. Given the very large blocks of alternative ancestry found in across all chromosomes, and that all sampled members of the KM population share the admixed regions, we speculate that the KM lineage may have been formed by a hybridisation event that was followed by very low levels of backcrossing with its parent populations.

## Discussion

Here, we have used comparative genomics and population genetics to dissect the evolutionary history of the English and Indian grain aphids, *Sitobion avenae* and *Sitobion miscanthi* – two important pests of cereal crops. It has been debated whether these species should be considered separate species (reviewed in Choe *et al*. 2006). The high-quality chromosome-scale genome assemblies generated here and our analysis of sequence divergence patterns revealed that *S. avenae* and *S. miscanthi* have less than 1% genome-wide sequence divergence (**Figure 3**), and hence, that they are closely related. Surprisingly, we found that the *S. miscanthi* isolate used for genome assembly is likely an F1 hybrid, with one of its haplotypes being more closely related to *S. avenae* than the other (**Figure 4**). Rather than being a two species system, *S. avenae* and *S. miscanthi* appear to belong to a species complex with multiple highly differentiated lineages that predate modern agriculture, having diverged between 199 thousand and 12 thousand years ago based on coalescent analysis (**Figure 5**). The species flock lineages have a highly reticulated evolutionary history (**Figure 6**) with evidence of hybridization, particularly among three Chinese lineages where we find large blocks of chromosomes with alternative ancestry shared by all members of each lineage (**Figure 7)**. We hypothesise that hybrid speciation is responsible for the evolution of new lineages in the *Sitobion* genus.

The unusual reproductive mode employed by aphids – cyclical parthenogenesis – may enable occasional high fitness combinations of parental lineages to rapidly expand through clonal propagation, effectively freezing a mosaic genome architecture as observed in Chinese *S. miscanthi*-like populations in this study. By avoiding or minimising sexual recombination after the hybridisation event, the descendants of hybrid aphids do not suffer from hybrid-vigour breakdown. Such breakdown leads to a fitness loss in sexual descendants, which is caused by segregation of the initially heterozygous loci and the breakdown of positive epistatic interactions (Moulia *et al*. 1995; Burke and Arnold 2001). Therefore, aphids may capitalise on the initial advantage of hybridisation (increasing variation and masking the genetic load, [Bertorelle *et al*. 2022]), whilst their ability to proliferate asexually after the event reduces future fitness costs. These factors may increase the likelihood of hybrid lineage formation in aphids. Indeed, the clonal proliferation of “frozen hybrid” lineages may be pervasive across aphid evolution as signatures of ancient hybridisation have been recently detected in a phylogenomic analysis of Aphididae (Owen and Miller 2022).

Finally, the genetic exchange between previously isolated lineages may play an important role in biological invasions of pests in our increasingly globalised world (Stukenbrock 2016; Rogério *et al*. 2022; Sotiropoulos *et al*. 2022). The benefits that hybridisation offers is preserved in clonal lineages, which gives clonal reproduction in facultative sexuals two important advantages. First, clonal reproduction increases colonising ability of pest species by not being reliant on conspecifics for reproduction. Second, hybridisation between diverged lineages generates novel genotypic variation that is important in adaptive evolution. Both advantages are currently capitalised on by many pest species because they can take advantage of human-mediated transport to colonise new habitats and host species. In addition, given that the hosts in agriculture tend to have little genetic diversity, a single successful genotype of a parasite or pest could infect an entire crop (Stukenbrock and McDonald 2008; Inoue *et al*. 2017; Latorre *et al*. 2020). However, our current study shows that this is not an evolutionary scenario unique to modern times, but that aphids may have exploited these evolutionary advantages of high mobility and cyclical parthenogenesis long before the advent of agriculture.

## Methods

### Genome assembly approach and quality control

For this study we generated *de novo* genome assemblies of *S. miscanthi, S. avenae, M. dirhodum* and *R. padi* using a variety of sequencing approaches detailed in the sections below. Regardless of the method used, we aimed to generate high-quality haploid genome assemblies maximising assembly completeness and minimising the inclusion of erroneously duplicated content (i.e. haplotigs). Genome assemblies were assessed by generating K-mer spectra, a procedure that involved comparing K-mer content of the raw sequencing reads to the K-mer content of the genome assembly with the K-mer analysis toolkit (KAT; [Mapleson *et al*. 2017]). We also assessed assembly completeness and duplication levels by searching for arthropod Benchmarking sets of Universal Single-Copy Orthologs (BUSCOs; n=1,066) using BUSCO v3 (Simão *et al*. 2015; Waterhouse *et al*. 2018). BUSCO and K-mer spectra analyses were used throughout the assembly process and to assess the final frozen genome assembly of each species. Each *de novo* assembly was checked for contamination and the presence of symbiont genomes by generating taxon-annotated GC content-coverage plots (known as a “BlobPlots”) with BlobTools v1.0.1 (Kumar *et al*. 2013; Laetsch and Blaxter 2017). Where symbiont genomes were co-assembled with their aphid host, we have included them as separate assembly files as part of the data release for this study. However, symbiont genomes have not been subjected to further curation or quality control.

### S. miscanthi *v1 genome evaluation*

We assessed the quality of the previously published *S. miscanthi* genome (Simis_v1; [Jiang *et al*. 2019]) using BUSCO and by generating a K-mer spectra comparing the published Illumina short-reads (NCBI accession number: SRX5767526) to the genome assembly. To check synteny with the closely related species *A. pisum*, we aligned Simis_v1 chromosome-scale scaffolds to chromosome-scale scaffolds from the *A. pisum* JIC1 v1 assembly (Mathers *et al*. 2021) using the D-GENIES server (Cabanettes and Klopp 2018). To assess scaffolding quality, we visualised Hi-C contacts across the published genome assembly using Juicebox Assembly Tools (JBAT; Dudchenko *et al*. 2018). Hi-C reads (SRX5767527) from Jiang *et al*. (2019) were mapped to Simis_v1 using Juicer (Durand *et al*. 2016) with default settings and the resulting merged_nodups.txt file was processed using the run-assembly-visualizer.sh script from the 3dDNA assembly pipeline (Dudchenko *et al*. 2017).

### Reassembly of S. miscanthi

We reassembled the *S. miscanthi* genome using sequence data from Jiang *et al*. (2019). These data include PacBio long-reads (SRX5767529; 85x coverage), Illumina short-reads (SRX5767526; 105x coverage) and *in vivo* Hi-C data (SRX5767527; 76x coverage). *De novo* assemblies of the PacBio long reads were generated with Flye v2.8.1 (Kolmogorov *et al*. 2019) using default PacBio parameters (“--pacbio-raw”) and wtdgb2 v2.3 (Ruan and Li 2019) with default parameters. The wtdgb2 assembly was subjected to a single round of long-read polishing with wtpoa-cns using minimap v2.14 (Li 2018) PacBio read alignments with the parameter “-ax map-pb”. The Flye and wtdgb2 assemblies were merged using quickmerge v0.3 (Chakraborty *et al*. 2016) with the parameters “-l 1256119 -ml 10000 [Flye_assembly_fasta] [wtdgb2_assembly_fasta]”. The “-l” flag was conservatively set to the N50 of wtdbg2 assembly as the Flye assembly had an N50 below 1 Mb (scaffold N50 = 583 Kb) and low values of “-l” may lead to increased misjoins. The Flye assembly was used as the “query” sequence because preliminary analysis showed it to be more complete than the wtdgb2 assembly and quickmerge assemblies predominantly contain sequence content from the “query” assembly. The quickmerge assembly was subjected to a single round of long-read polishing using the Flye polisher and followed by three rounds of short-read polishing with Pilon v1.22 (Walker *et al*. 2014). Redundant haplotigs were removed from the polished assembly using purge_dups (Guan *et al*. 2020). For purge_dups, scaffold coverage was estimated by mapping the PacBio long reads with mininmap v2.16 with the parameter “-x map-pb” and assembly self-alignment was carried out with minimap v2.16 with the parameter “-xasm5 -DP”. Scaffold coverage cutoffs for purge_dups were estimated automatically using the calcuts script.

We scaffolded the draft assembly into chromosome-scale super scaffolds using the published Hi-C data (Jiang *et al*. 2019). The Juicer pipeline was used to identify Hi-C contacts and the 3D-DNA assembly pipeline was used for assembly scaffolding (with default parameters), followed by manual curation with JBAT. We found that the Hi-C library had low resolution, resulting in sub-optimal scaffolding performance by 3D-DNA. However, 3D-DNA first orders the input assembly into a single super-scaffold before breaking the assembly into putative chromosome-scale fragments. Inspection of the initial round of scaffolding revealed sufficient signal to manually assemble the *S. miscanthi* chromosomes in JBAT (**Supplementary Figure 16**). The scaffolded assembly was screened for contamination based on manual inspection of “BlobPlots”. Briefly, short-reads were aligned to the assembly with BWA mem v0.7.7 (Li 2013) and used to estimate average coverage per scaffold. Additionally, each scaffold in the assembly was compared to the NCBI nucleotide database (nt; downloaded 13^th^ October 2017) with BLASTN v2.2.31 (Camacho et al. 2009) with the parameters “-task megablast - culling_limit 5 -evalue 1e-25 -outfmt ‘6 qseqid staxids bitscore std sscinames sskingdoms stitle’. Read mappings and blast results were passed to BlobTools v1.0.1 which was used to generate “BlobPlots” annotated at the order and genus level (**Supplementary Figures 17** and **18**). We removed two scaffolds belonging to the obligate *Buchnera* endosymbiont and additional scaffolds that had low coverage (<30x Illumina short-read converge). The remaining scaffolds were ordered by size and assigned a numbered scaffold ID with SeqKit v0.9.1 (Shen *et al*. 2016) to create a frozen release for downstream analysis (Simis_v2).

### Sequencing and de novo assembly of S. avenae JIC1 and M. dirhodum

*S. avenae* and *M. dirhodum* individuals were sampled from clonal lineages maintained at the JIC insectary on *Avena sativa* (oats). The *S. avenae* colony (dubbed JIC1) was originally obtained from the University of Newcastle in 2012. The original plant host is unknown. The *M. dirhodum* colony (dubbed UK035) was originally collected from a rose bush in Norwich in 2015.

We followed the procedures described in Mathers *et al*. (2020) to generate low-cost short-read *de novo* genome assemblies of *S. avenae* and *M. dirhodum*. Briefly, DNA was extracted from a single individual and sent to Novogene (China) where a PCR-free Illumina sequencing library was prepared with a target insert size of 500 – 1000 bp and sequenced on an Illumina HiSeq 2500 instrument with 250 bp paired-end chemistry. We also extracted total RNA from bulked adult un-winged asexual female individuals from each species and sent it to Novogene for strand-specific library preparation and sequencing on an Illumina platform with 150 bp paired-end chemistry. Genomic reads were processed with trim_galore (http://www.bioinformatics.babraham.ac.uk/projects/trim_galore) to remove adapters with the parameters “--quality --paired --length 150” and then assembled using Discovar *de novo* (https://software.broadinstitute.org/software/discovar/blog/) with default parameters. Erroneously duplicated content (i.e. haplotigs) in the initial draft assemblies was identified and removed using the K-mer based deduplication pipeline described in Mathers *et al*. (2020). Following deduplication, the assemblies were screened for contamination based on manual inspection of “BlobPlots” generated as described above for *S. miscanthi* (*S. avenae*: **Supplementary Figures 19** and **20**; *M. dirhodum*: **Supplementary Figures 21** and **22**). For *S. aveane*, we identified and removed three circular scaffolds corresponding to the chromosome (636 Kb long) and two plasmids of the obligate endosymbiont *Buchnera aphidicola*. We also removed additional scaffolds that had low coverage (< 15x Illumina short-read converge). For *M. dirhodum*, we identified and removed a circular scaffold corresponding to the *Buchnera* chromosome (642 Kb long) and three scaffolds (two circular) corresponding to *Buchnera* plasmids. We also identified and removed 121 scaffolds corresponding to the secondary symbiont *Regiella insecticola*. The *R. insecticola* scaffolds spanned 2.8 Mb which is similar to the reported genome size of an *R. insecticola* isolate from pea aphid (Nikoh *et al*. 2020), indicating we have likely assembled the complete genome of this bacterium. We also removed additional scaffolds that had low coverage (< 10x Illumina short-read converge).

We further improved the contiguity of the *S. avenae* and *M. dirhodum* assemblies using RNA-seq scaffolding with P_RNA_scaffolder (Zhu *et al*. 2018) as described in Mathers *et al*. (2020). For *S. avenae*, the RNA-seq scaffolded assembly was carried forward for further scaffolding with Hi-C data (see section below). For *M. dirhodum*, an additional round of assembly deduplication was carried out using purge_dups with assembly self-alignment carried out as for *S. miscanthi* and scaffold coverage estimated by mapping the PCR-free Illumina library to the draft assembly with BWA mem v0.7.7 with default parameters. Coverage cut-offs for purge_dups were set manually with the calcuts script with the parameters “-l 5 -m 25 -u 90 “. Finally, the *M. dirhodum* assembly was ordered by size and assigned a numbered scaffold ID with SeqKit v0.9.1 to create a frozen release for downstream analysis (Medir_v1.1).

### Sequencing and de novo assembly of R. padi

*R. padi* individuals were sampled from a clonal lineage (dubbed JIC1) maintained at the JIC insectary on *A. sativa* (oats). The colony was originally sampled in 2005. The original plant host and sampling location is unknown.

We followed procedures described in Biello *et al*. (2021) to extract high molecular weight DNA from a single *R. padi* individual and sent this to Novogene for 10x genomics link-read sequencing (Zheng *et al*. 2016). We generated an initial *de novo* assembly with Supernova v2.1.1 (Weisenfeld *et al*. 2017) with the parameter “--maxreads=143797524” set to give approximately 56x coverage. To increase contiguity of the Supernova assembly we carried out two rounds of linked-read scaffolding with scaff10x (https://github.com/wtsi-hpag/Scaff10X) with the parameters “-longread 0 -edge 45000 -block 45000” followed by a single round of misjoin detection and scaffolding with Tigmint v1.1.2 (Jackman *et al*. 2018) using default parameters. The resulting draft assembly was carried forward for further scaffolding with Hi-C data (see section below).

### S. avenae and R. padi multi-species Hi-C library preparation and genome scaffolding

Whole bodies of *R. padi* and *S. avenae* individuals from clonally reproducing colonies maintained at the JIC Insectary were snap frozen in liquid nitrogen, pooled in approximately equal numbers (∼100 aphids in total) and sent to Dovetail Genomics (Santa Cruz, CA) for Hi-C library preparation and sequencing on an Illumina HiSeq X instrument with 150 bp paired-end chemistry. Hi-C library preparation was carried out using the DpnII restriction enzyme following a similar protocol to Lieberman-Aiden *et al*. (2009). We scaffolded both assemblies using the same multi-species Hi-C library with the 3dDNA assembly pipeline (with default settings) followed by manual curation with JBAT. Pre curation Hi-C contact maps are shown in **Supplementary Figures 23** and **24** for *S. avaenae* and *R. padi*, respectively.

For *R. padi*, we screened the resulting chromosome-scale assembly for contamination based on manual inspection of “BlobPlots” generated as described above for *S. miscanthi* (**Supplementary Figures 25** and **26**) and identified a fragmented assembly (159 short scaffolds) of the obligate endosymbiont *Buchnera* which was removed from the final assembly. We also removed additional low coverage scaffolds (< 20x linked-read coverage). For *S. avenae*, following manual curation of the 3dDNA assembly, we removed additional duplicated content (haplotigs) from the assembly with purge_dups with scaffold coverage estimated from mapping the PCR-free Illumina library with BWA mem v0.7.7 and assembly self-alignment with minimap v2.16 with the parameters “-xasm5 -DP”. Coverage cutoffs for purge_dups were estimated automatically using the calcuts script. Finally, the *S. avenae and R. padi* genome assemblies were ordered by size and assigned a numbered scaffold ID with SeqKit v0.9.1 to create a frozen releases for downstream analysis (*S. avenae*: Siave_v2.1; *R. padi*: Rhpad_v1).

### Genome annotation

Our new assemblies of *S. miscanthi, S. avenae, M. dirhodum* and *R. padi* were annotated following Mathers *et al*. (2020b) incorporating evidence from RNA-seq data. Each genome was soft-masked with RepeatMasker v4.0.7 (Tarailo-Graovac and Chen 2009; Smit *et al*. 2015) using known Insecta repeats from Repbase (Bao *et al*. 2015) with the parameters “-e ncbi - species insecta -a -xsmall -gff”. RNA-seq reads were mapped to the genomes with HISAT2 v2.0.5 (Kim *et al*. 2015) with the parameters “--max-intronlen 25000 --dta-cufflinks” followed by sorting and indexing with SAMtools v1.3 (Li *et al*. 2009). Where strand-specific RNA-seq reads were available, we included the parameter “--rna-strandness RF”. We then ran BRAKER2 (Hoff *et al*. 2015, 2019) with UTR training and prediction enabled with the parameters “--softmasking --gff3 --UTR=on”. Strand-specific RNA-seq alignments were split by forward and reverse strands and passed to BRAKER2 as separate BAM files to improve the accuracy of UTR models as recommended in the BRAKER2 documentation. For *S. miscanthi*, we used unstranded RNA-seq data from Jiang *et al*. (2019). For *S. avenae* and *M. dirohdum* we used strand-specific RNA-seq generated for this study derived from pools of unwinged adult asexual females. For *R. padi*, we used un-stranded RNA-seq data from Thorpe *et al*. (2016). Full details of RNA-seq libraries used for genome annotation are given in **Supplementary Table 1**. Following gene prediction, genes were removed that contained in frame stop codons using the BRAKER2 script getAnnoFastaFromJoingenes.py and the completeness of each gene set was checked with BUSCO v3 with the Arthropoda gene set (n=1,066), using the longest transcript of each gene as the representative transcript. For *S. miscanthi*, we compared RNA-seq pseudo alignment rates between the published v1 annotation from Jiang *et al*. (2019) and our new annotation based on the Simis_v2 assembly. The *S. miscanthi* RNA-seq library used for both annotations was pseudo aligned to the Simis_v1 and Simis_v2 transcript sets with Kallisto v0.44.0 (Bray *et al*. 2016) with 100 bootstrap replicates (all other parameters were default) and alignment rates were extracted from the Kallisto run reports.

### Phylogenomic analysis

Protein sequences from our new genome assemblies of *S. miscanthi, S. avenae* and *M. dirhodum* and eight previously published Aphidinae genomes (Nicholson *et al*. 2015; Thorpe *et al*. 2018; Chen *et al*. 2019; Mathers 2020; Mathers *et al*. 2020, 2021) were clustered into orthogroups with OrthoFinder version 2.3.8 (Emms and Kelly 2015, 2019). Genome assembly and annotation versions are summarised in **Supplementary Table 3**. Where genes had multiple annotated transcripts, we used the longest transcript to represent the gene model. OrthoFinder was run in multiple sequence alignment mode (“-M msa -S diamond -T fasttree”) with DIAMOND version 0.9.14 (Buchfink *et al*. 2014), Multiple Alignment using Fast Fourier Transform (MAFFT) version 7.305 (Katoh and Standley 2013) and FastTree version 2.1.7 (Price, Dehal, & Arkin, 2009, 2010) used for protein similarity searches, multiple sequence alignment and gene and species tree estimation, respectively. The OrthoFinder species tree was automatically rooted based on informative gene duplications with Species Tree Root Inference from Gene Duplication Events (STRIDE; Emms and Kelly 2017). For visualisation, the species tree was pruned to only include dactynotine aphids (*S. miscanthi, S. avenae, M. dirhodum* and *A. pisum)* and *M. persicae* (outgroup) using ape v5.1 (Paradis and Schliep 2019). Genome size, gene counts and orthology information were visualised on the phylogeny with Evolview v3 (Subramanian *et al*. 2019).

### Synteny analysis

We identified syntenic blocks of genes between *S. avenae* (Simis_v2.1) and *S. miscanthi* (Simis_v2), and the published chromosome-scale genome assemblies of *A. pisum* (JIC1 v1) and *M. persicae* (clone O v2) (Mathers *et al*. 2021) using MCScanX v1.1 (Wang *et al*. 2012). For each comparison, we carried out an all versus all BLAST search of annotated protein sequences using BLASTALL v2.2.22 (Altschul *et al*. 1990) with the parameters “-p BlastP - e 1e-10 -b 5 -v 5 -m 8” and ran MCScanX with the parameters “-s 5 -b 2,” requiring synteny blocks to contain at least five consecutive genes and to have a gap of no more than 20 genes. MCScanX results were visualized with SynVisio (Bandi and Gutwin 2020).

### Whole genome alignment and estimation of sequence divergence

We used Progressive Cactus v1.0.0 (Armstrong *et al*. 2020) to align *S. avenae* v2.1 (Simis), *S. miscanthi* v2 (Simis), *A. pisum* JIC1 v1 (Acpis) and *M. persicae* clone O v2 (Myper) genomes given the phylogeny (((Siave,Simis),Medir),Acpis) with default parameters. Tools from the hal (Hickey *et al*. 2013) and PHAST v1.5 (Hubisz *et al*. 2011) packages were used to manipulate the alignment and calculate divergence statistics. Alignment coverage statistics relative to *S. avenae* and *S. miscanthi* were calculated using halStats with the “--coverage” option. To carry out window-based pairwise sequence divergence analysis relative to the *S. miscanthi* v2 reference genome, we specified fixed 1 Mb windows along *S. miscanthi* chromosome-scale scaffolds using makewindows from bedtools v2.28.0 (Quinlan and Hall 2010) and extracted alignments for each window in maf format using hal2maf with the parameters “--refGenome Simis --noAncestors --onlyOrthologs --refTargets [window bed file]”. The maf files were post processed with maf_stream merge_dups (https://github.com/joelarmstrong/maf_stream) in “consensus” mode to resolve alignments to multiple genomic copies as described Feng *et al*. (2020). The maf_stream processed alignment files were converted to fasta format with msa_view with the parameter “--soft-masked”. To generate pairwise divergence estimates for each genomic window, we reduced the fasta formatted alignment files to contain sequences from *S. miscanthi* and one other target species (either *S. avenae, M. dirhodum* or *A. pisum*) with SeqKit v0.9.1 (seqkit grep) and passed these files to PhlyoFit which was run with default settings to estimate divergence in substitutions per site under the REV model.

### JIC1 and Langfang-1 haplotype divergence

To investigate intra- and inter-individual patterns of sequence divergence in *S. miscanthi* and *S. avenae* we reconstructed independent phased haplotypes for each assembly using HapCUT2 v1.1 (Edge *et al*. 2017) and generated a four-way whole genome alignment with sibeliaZ (Minkin and Medvedev 2020). Our approach took advantage of the availability of *in vivo* Hi-C data for both isolates, which contains accurate long range phasing information (Edge *et al*. 2017), and the unique biology of aphids which means lab reared colonies can be maintained as clonal lineages in the absence of recombination (aphid parthenogenesis is apomictic [Blackman 1979; Tomiuk and Wöhrmann 1982; Hales *et al*. 2002]). As such, although sequence data for each isolate is derived from pools of individuals (except from PCR-free Illumina sequence data for *S. avenae*), all individuals sequenced for a given isolate are expected to contain the same two haplotypes and so sequence data can be combined to reconstruct fully phased haplotypes for each isolate.

We followed the HapCUT2 pipeline to assemble chromosome-scale haplotypes for *S. aveane* JIC1 and *S. miscanthi* Langfang-1. Short-read data for each isolate was mapped to its respective reference genome with BWA mem v0.7.17 and the resulting alignments were sorted and indexed with SAMtools v1.7 followed by PCR duplicate marking with picard MarkDuplicates v2.1.1 (https://broadinstitute.github.io/picard/). Using these data, we called single nucleotide polymorphisms (SNPs) and short structural variants (indels) with Freebayes v1.3.1 (Garrison and Marth 2012) with default parameters. The initial variant sets were filtered with BCFtools v1.8 (Danecek *et al*. 2021) and VcfFilter (https://github.com/biopet/vcffilter) to retain biallelic sites and remove low quality sites (QUAL < 30) and sites with low sequence coverage (DP < 5). The filtered variant file (in vcf format) was then split by chromosome using BCFtools view for processing with HapCUT2. Next, we extracted haplotype informative information from our read sets for each chromosome using extractHAIRS from HapCUT2. For *S. miscanthi*, we used phase information from PacBio long reads and *in vivo* Hi-C data. PacBio reads were aligned to Simis_v2 using minimap v2.14 with the parameter “-ax map-pb” and the resulting alignments sorted with SAMtools v1.9 and passed to extractHAIRS with the parameters “--pacbio 1 --new_format 1 --indels 1”. Hi-C reads were aligned separately for read 1 and read 2 using BWA mem v0.7.12 and the resulting alignments sorted by read name with sambamba (Tarasov *et al*. 2015) and passed to the HapCUT2 script HiC_repair.py to generate a merged alignment file. The repaired Hi-C mapping file was sorted by read name with sambamba, processed with SAMtools fixmate, sorted again by coordinate and PCR duplicates marked with picard MarkDuplicates v2.1.1. The processed Hi-C alignments were passed to extractHAIRS with the parameters “-- HiC 1 --new_format 1 --indels 1”. For *S. avenae*, in the absence of long-read data, we used phase information from our PCR-free reads and our *in vivo* Hi-C data. Preliminary analysis showed that including phase information from the PCR-free Illumina reads increased the proportion of phased variants on scaffold_1 (the longest chromosome in the assembly) from 79% (137,885 / 172,422) to 97% (166,986 / 172,422) compared to just using phase information from the Hi-C reads. *S. avenae* Hi-C reads were aligned to Siave_v2.1 and processed following the procedure described for *S. miscanthi*. For the *S. avenae* PCR-free Illumina reads, we used the alignment file generated for variant calling and passed it to extractHAIRS with the parameters “--new_fromat 1 --indels 1”. Using the variant calls and phase information generated by extractHAIRS we ran HapCUT2 separately for each chromosome of *S. miscanthi* and *S. aveane* with the parameters “--HiC 1 --ea 1 --nf 1 --outvcf 1”. Phasing statistics were extracted from the resulting vcf files with WhatsHap stats v0.17 (Martin *et al*. 2016) and we made fasta files of each haplotype (per chromosome) using BCFtools consensus v1.8 with haplotypes specified with either “-H 1” or “-H 2” and concatenated them by haplotype and sample (either *S. avaenae* JIC1 or *S. miscanthi* Langfang-1). Overall, this pipeline generated four independent haplotype assemblies (incorporating SNPs and indels) with >=99.96% of each chromosome contained in a single phase block (**Supplementary Table 5**). Although each chromosome is nearly fully phased for each sample, the assignment of H1 or H2 haplotype IDs is arbitrary between chromosomes.

To estimate divergence between the assembled haplotypes of *S. avenae* JIC1 and *S. miscanthi* Langfang-1 we generated a four-way whole genome alignment with sibeliaZ using default settings and processed the alignment with MafFilter v1.3.1 (Dutheil *et al*. 2014) and mafTools v0.2 (Earl *et al*. 2014). MafFilter subset with the parameters “species=(JIC1_H1,JIC1_H2,LF1_H1,LF1_H2),strict=yes,keep=no,remove_duplicates=yes)” was used to retain alignment blocks that are covered by all haplotypes and remove blocks containing paralogs. The filtered alignment was ordered using JIC1 haplotype 1 (JIC1_H1) as the reference with mafRowOrderer with the parameter “--order JIC1_H1,JIC1_H2,LF1_H1,LF1_H2” then processed with mafStrander and mafSorter, both with the parameter “--seq=JIC1_H1”. We specified 100 Kb fixed genomic windows relative to the JIC1_H1 assembly and estimated pairwise sequence divergence with phyloFit as for the divergence estimates generated for the progressive cactus alignment described in the section above, with the exception that mafExtractor was used to generate window specific alignment files from the pre-processed sibeliaZ maf file.

### Samples, read mapping and genotyping

We obtained genotyping be sequencing (GBS) data for *S. avenae* UK populations (119 samples) and *S. miscanthi* Chinese populations (100 samples) from Morales-Hojas *et. al*. (2020). Sample information is provided **Supplementary Table 6**. Reads were trimmed for adapters and low-quality bases with trim_galore v0.4.5 with the parameters “--paired --length 100”. We also included Illumina short-reads from the isolates of *S. avaenae* (JIC1) and *S. miscanthi* (Langfang-1) used for genome assembly. These data were sub-sampled to give approximately 25x coverage using SeqKit sample v0.9.1 with the parameters “-p 0.25 -s 1234” for *S. miscanthi* Langfang-1 and “-p 0.4 -s 1234” for of *S. avaenae* JIC1. All read sets were mapped to the *S. avenae* v2.1 genome assembly using BWA mem v0.7.17 with default parameters and the alignments were sorted and indexed with SAMtools v1.7 followed by PCR duplicate marking with picard MarkDuplicates v2.1.1. Mapping statistics were gathered with QualiMap v2.2.1 (Okonechnikov *et al*. 2016) and we omitted all samples with less than 1,000,0000 aligned reads (n = 12) from downstream analyses. Variant calling was carried out with Freebayes v1.3.1 with default parameters. Variants were filtered with BCFtools v1.8 as follows: we retained only biallelic SNP sites located on one of the nine chromosome-scale *S. avenae* v2 scaffolds, we removed sites with low quality (QUAL < 30) and individual genotype calls with fewer than two supporting reads (FORMAT/DP < 2). We then further filtered the variant set to remove sites with more than 25% missing data using VCFtools v0.1.15 (Danecek *et al*. 2011) with the parameter “--max-missing 0.75”. Following these steps, we identified and removed 60 samples that had more than 30% missing data (across all retained sites) using VCFtools v0.1.15.

### Principle component analysis

We investigated relationships among the Chinese and UK samples using principle component analysis with SNPrelate (Zheng *et al*. 2012). To minimise the effects of linkage disequilibrium (LD) SNPs were thinned to one SNP every 25 Kb with VCFtools v0.1.15 (Danecek *et al*. 2011), reducing the filtered variant set to 1,772 sites. Plotting of principle components 1 and 2 revealed clustering of the Chinese samples in accordance with the populations identified by Morales-Hojas *et. al*. (2020) which cluster largely based on geographic location. We therefore assigned each sample to either one of six Chinese populations identified by Morales-Hojas *et. al*. (2020) or to the UK population **Supplementary Table 6**. Seven samples (out of 149) clustered with a different population to that expected by their geographic origin – these samples were assigned to populations based their genetic identity (ascertained by PCA).

### Phylogenetic network analysis

To further visualise population structure in our data and to investigate the origin of the Langfang-1 individual used for *S. miscanthi* genome assembly we generated a distance-based split network using the neighbour-net algorithm with SplitsTree v4.14.6 (Huson and Bryant 2006). To generate the network, we phased the filtered variant set with BEAGLE v5.1 (Browning and Browning 2007) using default settings and thinned the phased SNP set to one SNP every 25 Kb with VCFtools v0.1.15. Haplotypes from *S. avenae* v2.1 scaffold_2 (the longest autosome) were extracted in fasta format using PGDspider v2.1.1.5 (Lischer and Excoffier 2012) with the parameters “FASTA_WRITER_HAPLOID_QUESTION=false VCF_PARSER_EXC_MISSING_LOCI_QUESTION=true VCF_PARSER_MONOMORPHIC_QUESTION=false VCF_PARSER_PLOIDY_QUESTION=DIPLOID”.

### Divergence time analysis

We used the pea aphid spontaneous mutation rate (2.7 × 10^−10^ per haploid genome per generation; [Fazalova *et al*. 2020]) to estimate the divergence time between *Sitobion* and *Metopolophium*. From our OrthoFinder phylogenomic analysis of aphids (see section above), we identified 11,702 phylogenetically inferred 1-to-1 orthologs between *S. miscanthi* and *M. dirhodum*. For each pair of orthologous genes, we extracted coding sequences, generated a codon alignment with PRANK v150803 (Löytynoja 2014) with the parameter “-codon” and estimated synonymous site (third codon positions) divergence using paml v4.9 (Yang 2007) with YN00 (Yang and Nielsen 2000). Genes were categorised based on their location (X chromosome or autosome) in the Simis_v2 genome assembly and genes on unplaced scaffolds were excluded. We then estimated the divergence time between *Sitobion* and *Metopolophium* (in number of generations) using the formula *T* = 0.5*d*_*S*_ / 2μ, where *d*_*S*_ is the median sequence divergence at autosomal synonymous sites and μ is the pea aphid spontaneous mutation rate in substitutions per haploid genome per generation. To convert our estimate from number of generations to years, we divided *T* by 15 which corresponds to the estimated number of aphid generations per year assuming 14 asexual generations and one sexual generation (Loxdale and Balog 2018).

To date the divergence of ingroup *Sitobion* lineages we jointly called variants among the *Sitobion* GBS samples (filtered set, see above), *S. miscanthi* Langfang-1 whole genome sample, *S. avaenae* JIC1 whole genome sample and the *M. dirhodum* whole genome sample and estimated a species / population tree under the MSC with SNAPP v1.5.1 (Bryant *et al*. 2012). To generate the variant set used for coalescent analysis, we mapped *M. dirhodium* PCR-free Illumina short reads to the *S. avaene* v2.1 genome assembly using BWA mem v0.7.17 with default parameters, sorted and indexed the alignments with SAMtools v1.7 and marked duplicates with picard MarkDuplicates v2.1.1. Variants among the *M. dirhodium* sample and the filtered set of *Sitobion* samples were called using Freebayes v1.3.1 with default parameters. Variants were filtered as for the *Sitobion*-only analysis with the exception that we removed sites with more than 10% missing data using VCFtools v0.1.15 with the parameter “--max-missing 0.9”. For the SNAPP analysis, we selected the two highest coverage samples from each population (**Supplementary Table 6**). Where populations contained samples from two locations, we selected the highest coverage sample from each location. The Langfang-1 sample was excluded due to putative hybrid origin. SNAPP xml files were prepared following Stange *et al*. (2018) using the script snapp_prep.rb. We set a starting tree that specified a split between *Sitobion* and *Metopolophium* with all other relationships unresolved (“(MedirOG:3.6,(UK:0.6,KM:0.6,LF_MY:0.6,SZ_PL:0.6,QD_TA:0.6,TG_YC:0.6,WH:0.6):3);”) and applied a normally distributed prior centred at 3.6 Mya (SD = 0.5 Mya) on the split between *Sitobion* and *Metopolophium* to calibrate the molecular clock. Ingroup *Sitobion* lineages were constrained to be monophyletic with respect to *M. dirhodum*. We carried out two independent SNAPP runs with BEAST v2.6.3 (Bouckaert *et al*. 2019), running each for 1 million MCMC iterations and taking samples every 500 iterations. We. checked stationarity and convergence of the runs with Tracer v1.7.1 (effective sample size > 100 for all parameters) and generated a maximum clade credibility tree using TreeAnnotator v2.6.3, discarding the first 10% of samples as burn in.

### Demographic history

We reconstructed historical changes in effective population size for *S. miscanthi* and *S. avaenae* using MSMC2 v2.0 (Malaspinas *et al*. 2016), which implements the multiple sequentially Markovian coalescent (MSMC) model. We used read mappings (against the *S. avenae* JIC1 v2.1 reference genome) from the population genomic analysis described above for the Langfang-1 and JIC1 whole genome samples and called variants in each sample with SAMtools mpileup v1.3 (parameters: “-q 20 -Q 20 -C 50 -u”) and BCFtools call v1.3.1 (parameters: “-c -V indels”). Variant calls from BCFtools were passed to the bamCaller.py script from the msmc-tools repository (github.com/stschiff/msmc-tools) to generate vcf and mask files which were in turn passed to the generate_multihetsep.py script (also from the msmc-tools repository) to generate the required input files for MSMC2. For bamCaller.py, we provided the average sequencing depth for the Langfang-1 (23x) and JIC1 (26x) samples as calculated from the output of SAMtools depth using only chromosome-scale scaffolds. MSMC2 was run with default setting and the output was scaled for plotting using the pea aphid spontaneous mutation rate (2.7 × 10^−10^ per haploid genome per generation; [Fazalova *et al*. 2020]) and 15 generations per year (Loxdale and Balog 2018).

### D statistics

We summarised admixture across the *S. avenae* / *miscanthi* complex using Patterson’s *D* (Patterson *et al*. 2012) and the *f*-branch (*f*_b_) statistic with Dsuite v0.4r38 (Malinsky *et al*. 2021). We used Dtrios from Dsuite with the SNP set generated for the SNAPP phylogenomic analysis (described above) and the SNAPP phylogeny to calculate *D*_min_, the minimum amount of allele sharing regardless of any assumptions made about the tree topology, for all trios of ingroup lineages (n=35). We also summarised rates of introgression using the *f*_b_ statistic with Fbranch from Dsuite using the “.tree” file generated by Dtrios and the SNAPP phylogeny. *f*_b_ statsitics were plotted on the SNAPP phylogeny using dtools.py, specifying *M. dirhodum* as the outgroup. We note that the Langfang-1 whole genome sample was excluded from this analysis due to putative hybrid origin.

### Topology weighting

To investigate genome wide patterns of introgression and hybridisartion among *Sitobion* lineages at greater resolution we re-filtered the raw variant calling results generated for the SNAPP phylogenomic analysis, excluding all the UK *S. avenae* GBS samples and the *M. dirhodum* sample. We expected the resulting set of filtered variants to contain more sites because we had previously implemented strict criteria requiring called sites to be shared by at least 90% of samples and the UK GBS samples had been prepared using a different restriction enzyme to the Chinese samples, meaning only a small number of sites overlapped in the two sets of GBS samples due chance proximity of restriction enzyme cut sites. We note that, in this reduced dataset, the UK *S. avenae* population is still represented by the JIC1 whole genome sample. After removing the UK GBS samples from the raw variant file we applied the following filtering criteria: we retained only biallelic SNP sites located on one of the nine chromosome-scale *S. avenae* v2 scaffolds, we removed sites with low quality (QUAL < 30) and individual genotype calls with fewer than two supporting reads (FORMAT/DP < 2), we removed sites with a called genotype in less than 90% of the samples. The filtered vcf file was then phased with BEAGLE v5.1 using default settings. We refer to this set of variants as the “large SNP set” in the section below.

We used topology weighting by iterative sampling of subtrees (Twisst; [Martin and Van Belleghem 2017]) to explore phylogenetic relationships across the genome, focussing on three focal lineages (WH, KM and QD_TA) inferred to have high levels of introgression based on the *f*_b_ analysis. We processed the phased “large SNP set” with scripts from the genomics_general repository (https://github.com/simonhmartin/genomics_general) to create phylogenetic trees (containing two haplotypes per sample) in 50 SNP windows across the *S. avenae* JIC1 v2.1 reference genome using PhyML v3.3 (Guindon *et al*. 2010) with the GTR substitution model. We then ran Twisst to calculate topology weightings for the three possible topologies describing relationships between the KM, WH, QD_TA and UK (included as an outgroup) lineages. Samples not belonging to the four lineages of interest were ignored. For visualisation of topology weightings along *S. avenae* JIC1 v2.1 chromosomes, a smoothing parameter was applied with a loess span of 1,000,000 bp, with a 100,000 bp spacing. SplitsTree networks were generated for the whole genome and for regions of interest as described above for the smaller SNP set.

## Data availability

Raw sequence data generated for this study are available at the NCBI short-read archive under BioProject PRJNA880698. Supporting data including OrthoFinder gene family clustering results, whole genome alignments, variant calls and SNAPP configuration files are available from Zenodo (https://doi.org/10.5281/zenodo.7108778). Genome assemblies and annotations generated in this study are available from the Aphidinae comparative genomics resource (https://doi.org/10.5281/zenodo.5908005).

## Acknowledgments

The described work was supported by a CEPAMS grant (17.03.2) awarded to S.H. and a Biotechnology and Biological Sciences Research Council (BBSRC) Future Leader Fellowship (BB/R01227X/1) awarded to T.C.M.. R.H.M.W. was funded from the BBSRC Norwich Research Park Biosciences Doctoral Training Partnership Award (BB/M011216/1). Additional support was provided by the BBSRC Institute Strategy Programs (BBS/E/J/000PR9797 and BBS/E/J/000PR9798) awarded to the John Innes Centre. The JIC is grant-aided by the John Innes Foundation. This research was supported by JIC Technology Platforms, particularly the Entomology and Insectary facilities and staff. We thank the NBI Computing Infrastructure for Science Group, which provides technical support and maintenance to the John Innes Centre’s high-performance computing cluster and storage systems.

