## Supplementary Figure for "Hybridisation has shaped a recent radiation of grass-feeding aphids"

**Supplementary Figures**

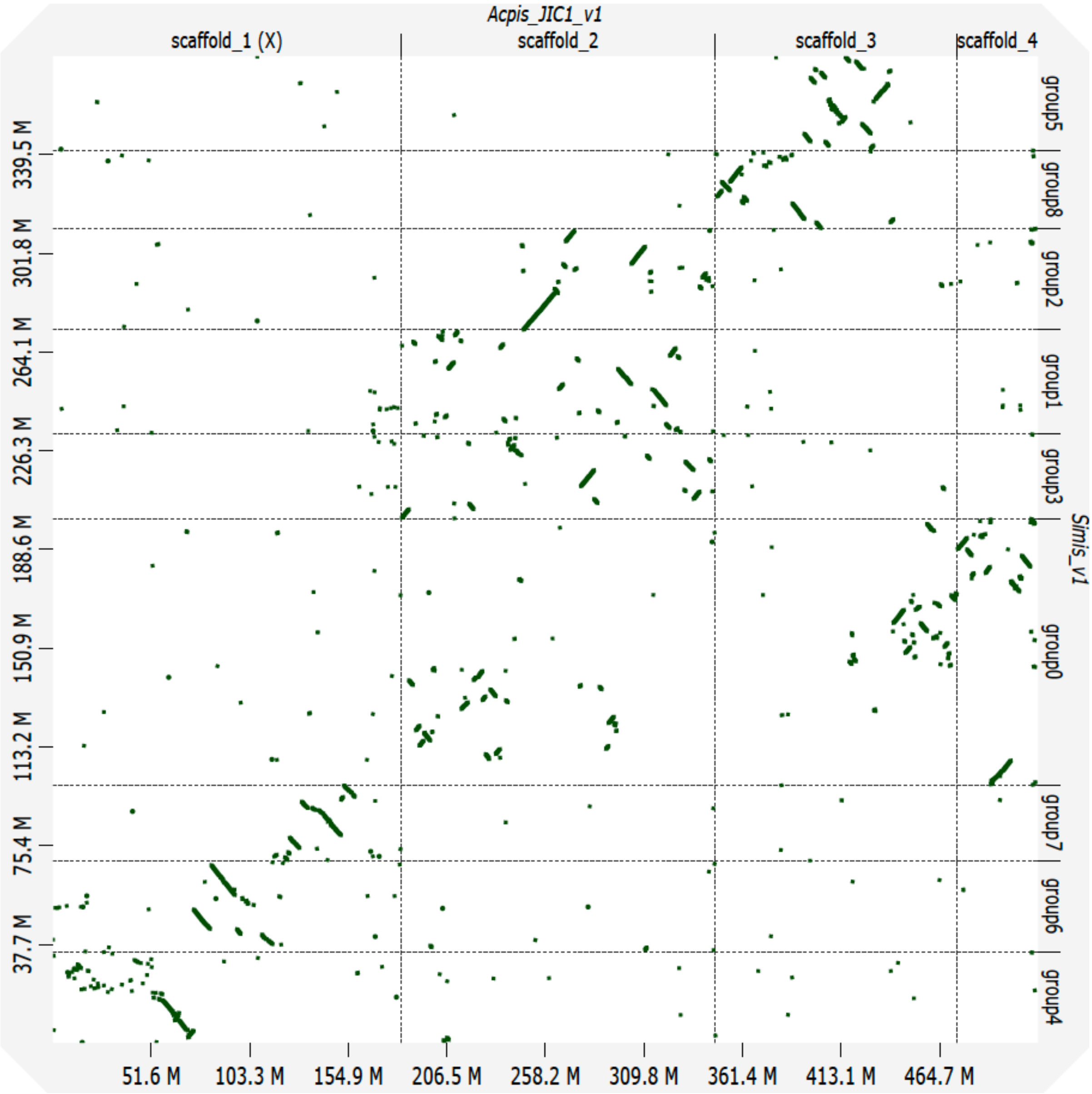


**Supplementary Figure 1:** *Dot plot* showing whole genome alignment of the *Sitobion miscanthi* v1 (y-axis; Simis_v1) and *Acyrthosiphon pisum* JIC1 v1 (x-axis; Acpis_JIC1_v1) genome assemblies. Simis_v1 scaffolds are ordered along the Acpis_JIC1_v1 assembly. For clarity, only chromosome scale genomic scaffolds are aligned. The x- and y-axis show cumulative scaffold length in Mb.

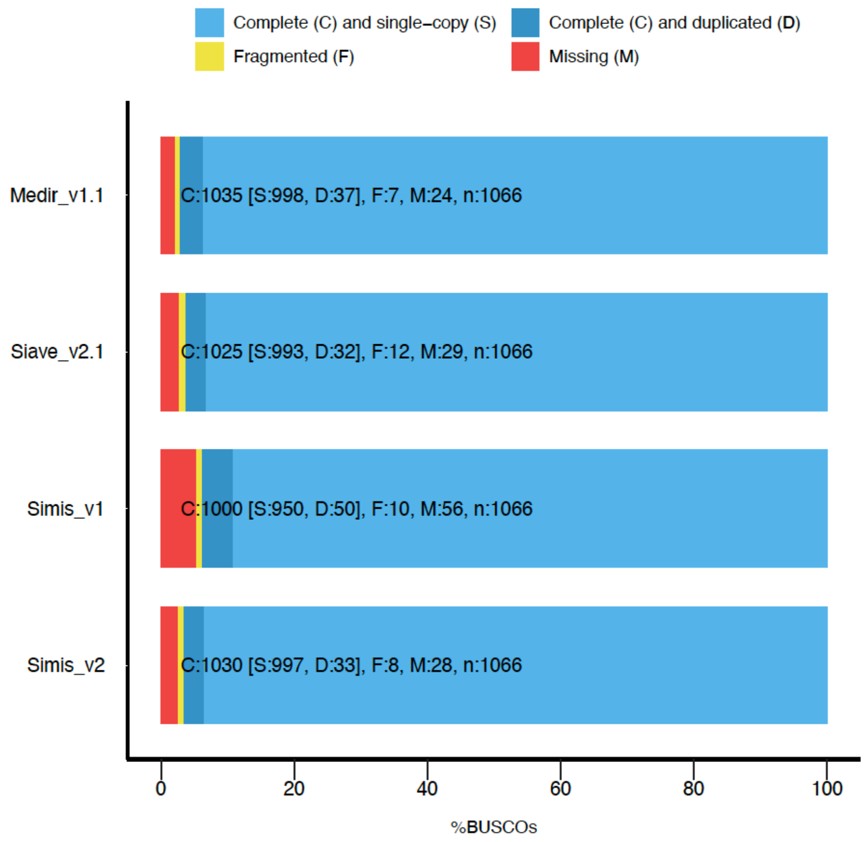


**Supplementary Figure 2:** BUSCO completeness plot for *Metopolophium dirhodum* v1.1 (Medir_v1.1), *Sitobion avenae* JIC1 v2.1 (Siave_v2.1), *Sitobion miscanthi* v1 (Simis_v1) and *Sitobion miscanthi* v2 (Simis_v2) genome assemblies. The genomes were assessed using BUSCO v3 and the Arthropoda gene set (odb9; n=1,066).

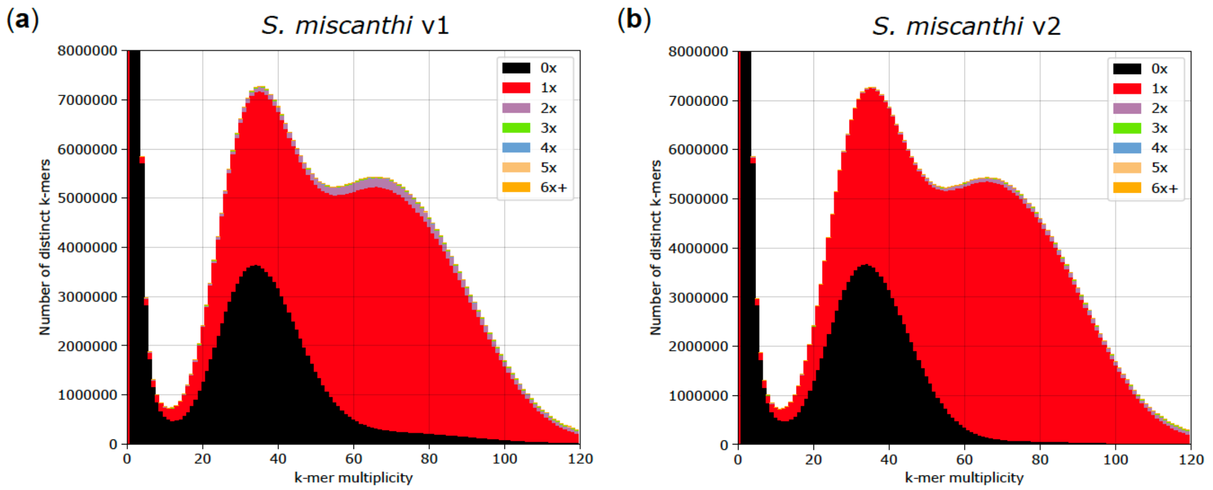


**Supplementary Figure 3:** KAT k-mer spectra plot comparing *S. miscanthi* Illumina paired-end (PE) reads from Jiang *et al.* (2019) to the original *S. miscanthi* assembly (**a**) and our updated *S. miscanthi* assembly (**b**). Colours indicate how many times fixed length words (k-mers) from the reads appear in the assembly. Red indicates k-mers found only once in the assembly, black indicates content present in the reads but missing from the assembly and other colours indicate k-mers that are duplicated in the assembly. The x-axis shows the number of times each k-mer is found in the reads (k-mer multiplicity) and the y-axis shows the count of distinct k-mers in 1x k-mer multiplicity bins.

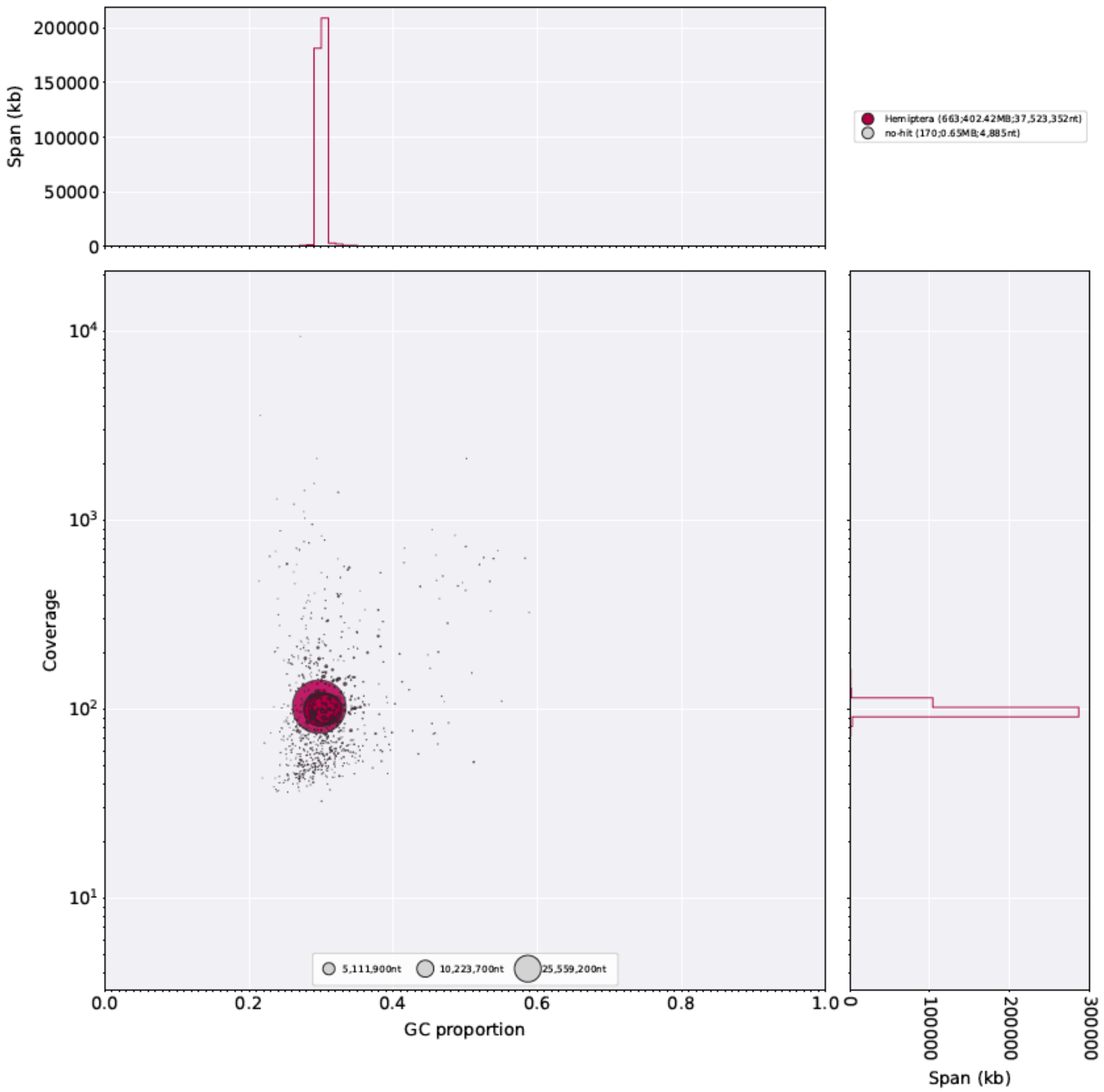


**Supplementary Figure 4:** Taxon-annotated GC content-coverage plot of the *Sitobion miscanthi* v2 genome assembly. Each circle represents a scaffold in the assembly, scaled by length, and coloured by order-level NCBI taxonomy assigned by BlobTools. The x-axis corresponds to the average GC content of each scaffold and the y-axis corresponds to the average coverage based on alignment with Illumina paired-end reads from the *S. miscanthi* Langfang-1 colony (from Jiang *et al.* 2019). Marginal histograms show cumulative genome content (in Kb) for bins of coverage (y-axis) and GC content (x-axis).

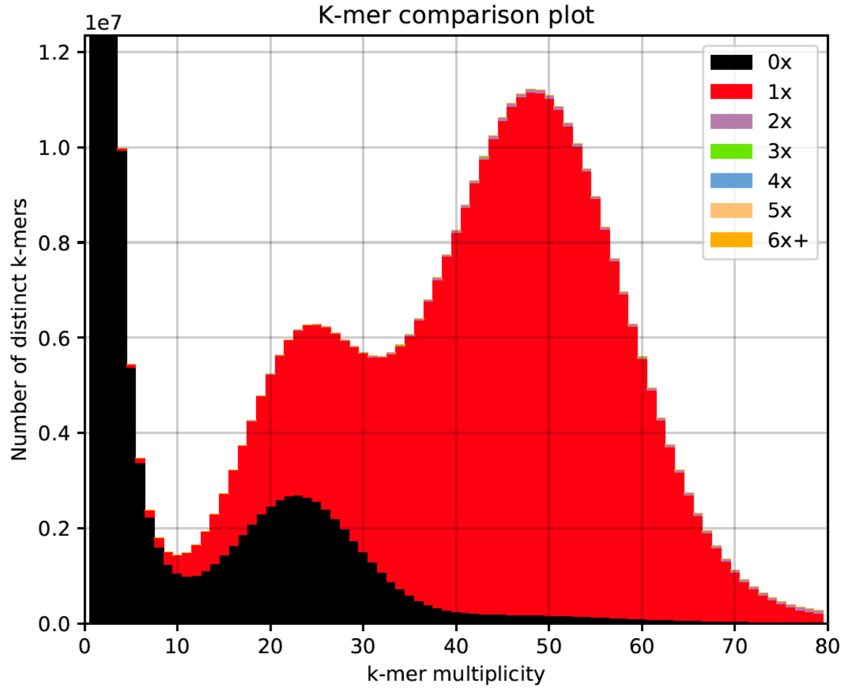


**Supplementary Figure 5:** KAT k-mer spectra plot comparing genomic *S. avenae* Illumina paired-end (PE) reads to the *S. avenae* v2.1 genome assembly. See **Supplementary Figure 3** legend for detailed description of the plot.

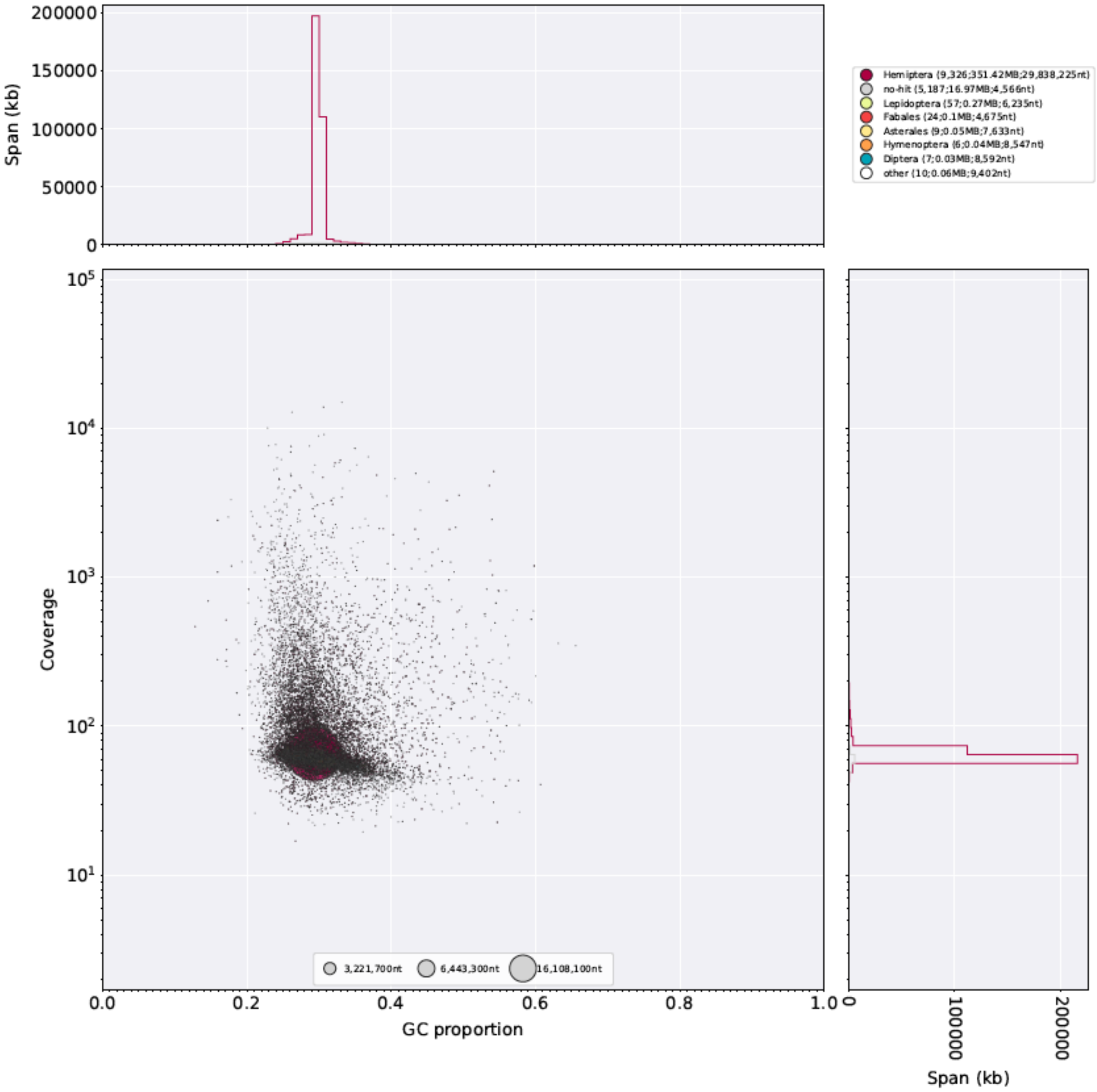


**Supplementary Figure 6:** Taxon-annotated GC content-coverage plot of the *Sitobion avenae* v2.1 genome assembly. Scaffold coverage (y-axis) is based on alignment of PCR-free Illumina paired-end reads from the *S. avenae* JIC insectary colony. See **Supplementary Figure 4** legend for detailed description of the plot.

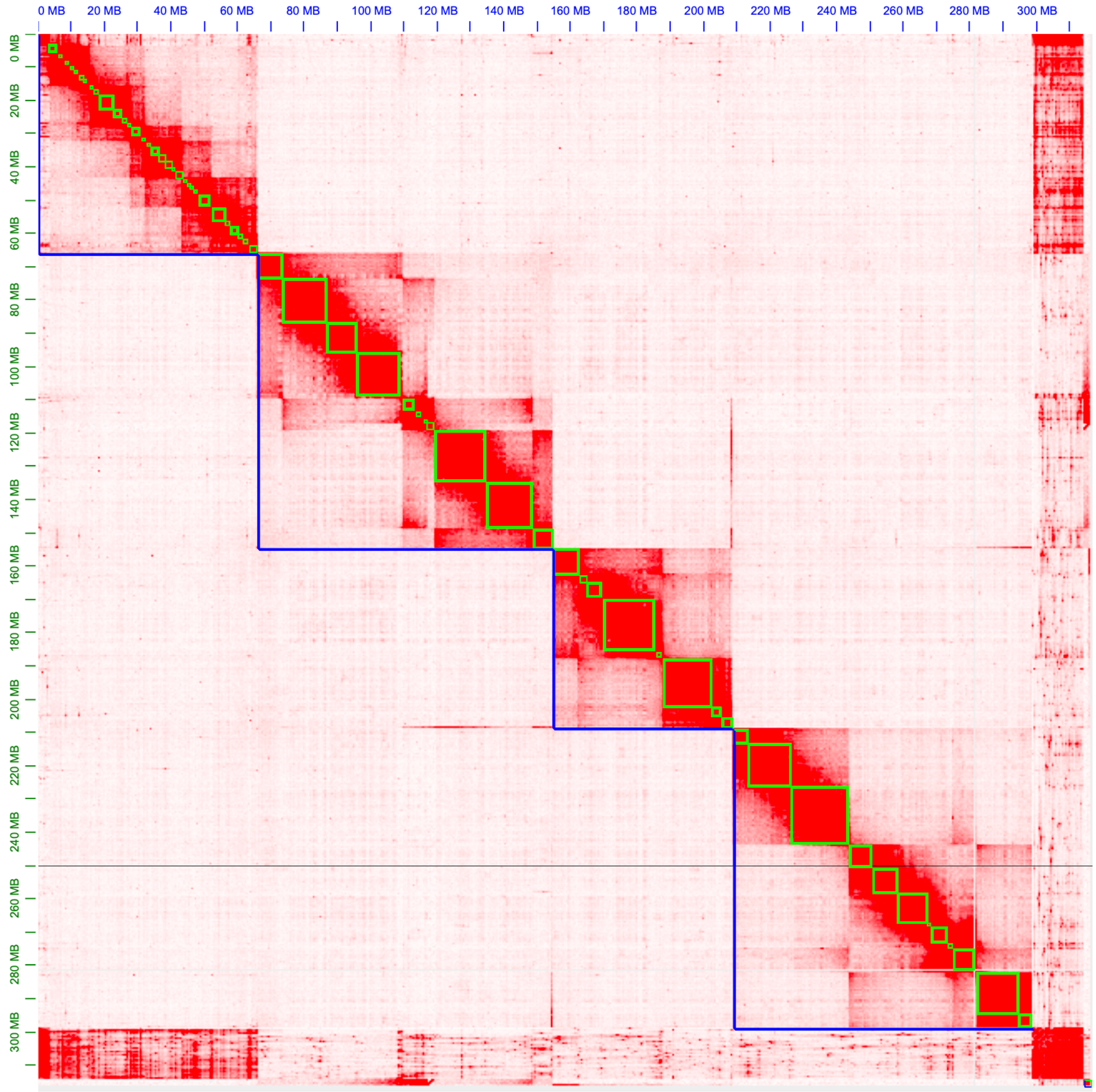


**Supplementary Figure 7:** Hi-C contact map for the *R. padi* v1 genome assembly. Blue lines show chromosome-scale super scaffolds, green lines show scaffolds from the 10x Genomics lined-read assembly. The x- and y-axis show cumulative length in millions of base pairs (Mb).

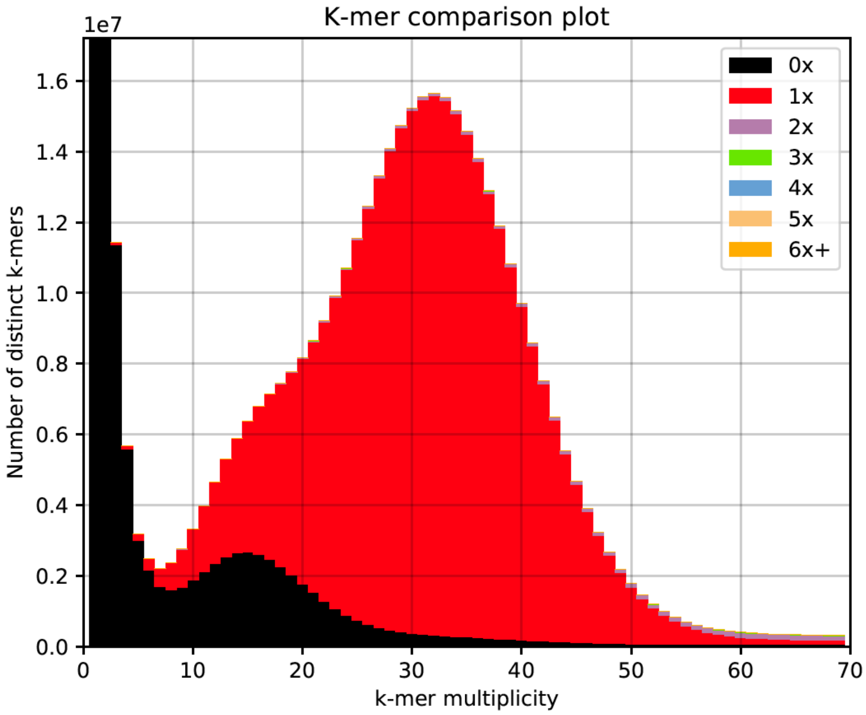


**Supplementary Figure 8:** KAT k-mer spectra plot comparing *Metopolophium dirhodum* Illumina paired-end (PE) reads to the *M. dirhodum* v1.1 genome assembly. See **Supplementary Figure 3** legend for detailed description of the plot.

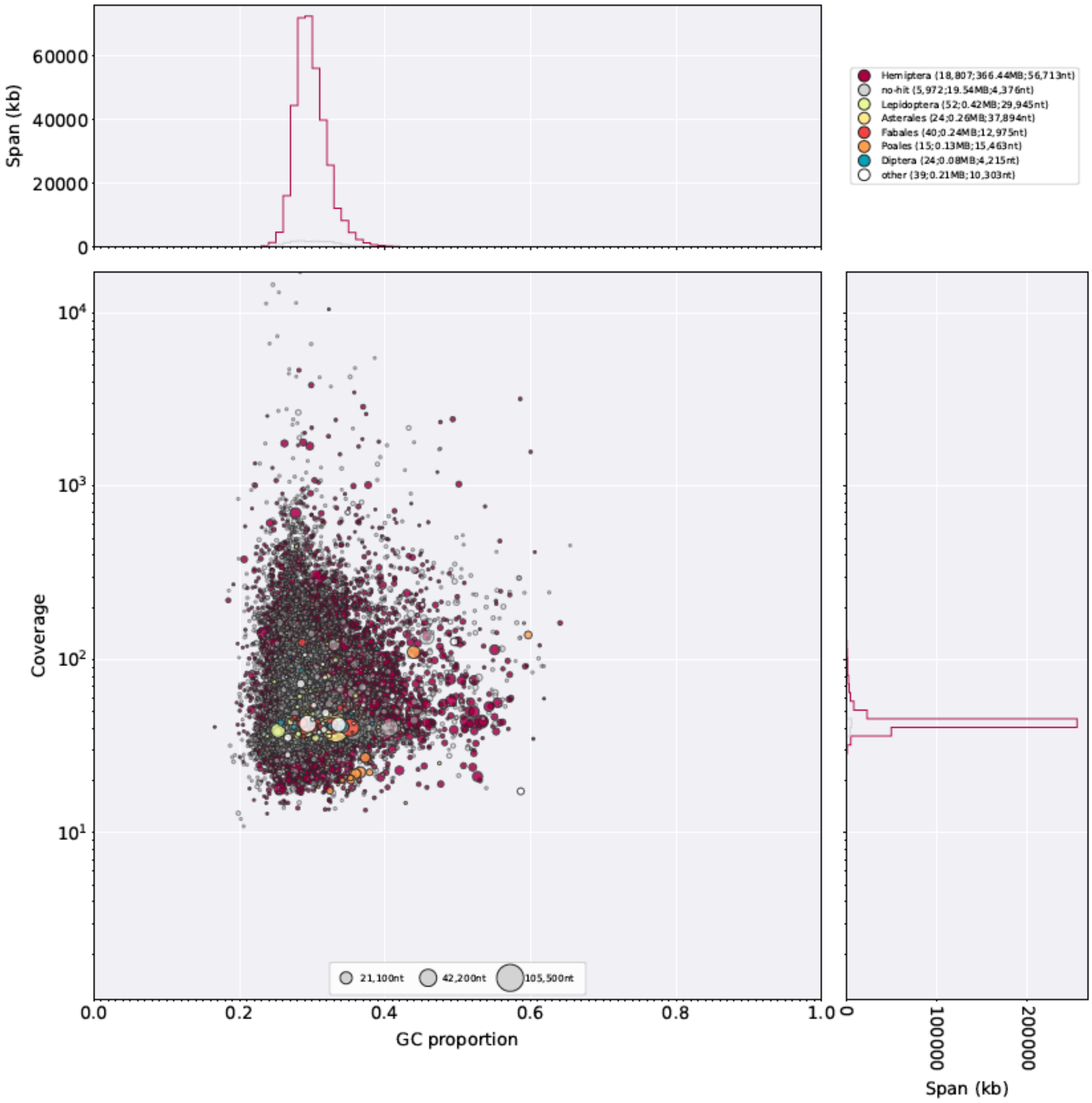


**Supplementary Figure 9:** Taxon-annotated GC content-coverage plot of the *Metopolophium dirhodum* v1.1 genome assembly. Scaffold coverage (y-axis) is based on alignment with PCR-free Illumina paired-end reads from the *M. dirhodum* JIC insectary colony. See **Supplementary Figure 4** legend for detailed description of the plot.

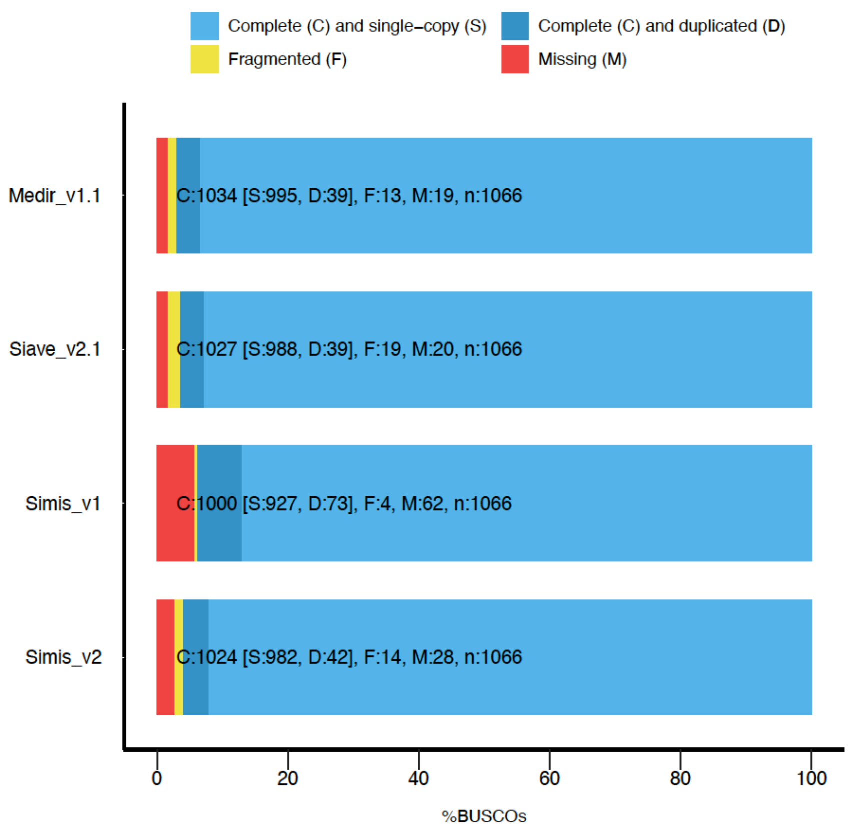


**Supplementary Figure 10:** BUSCO completeness plot for *Metopolophium dirhodum* v1.1 (Medir_v1.1), *Sitobion avenae* JIC1 v2.1 (Siave_v2.1), *Sitobion miscanthi* v1 (Simis_v1) and *Sitobion miscanthi* v2 (Simis_v2) gene sets. Proteomes from the annotation of each assembly were assessed using BUSCO v3 and the Arthropoda gene set (odb9; n = 1,066). Where multiple transcripts were annotated for a gene, we used the longest transcript.

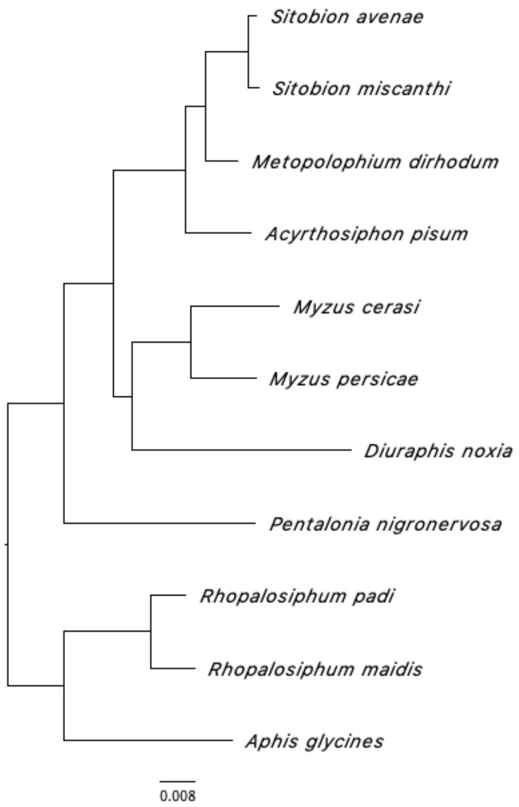


**Supplementary Figure 11:** Maximum likelihood phylogeny of 11 aphid species with sequenced genomes from the subfamily Aphidinae based on a concatenated alignment of 5,091 conserved one-to-one orthologues. The tree is rooted based on evidence from gene duplications with STRIDE. The basal node corresponds to the split between the tribes Macrosiphini and Aphidini. All nodes received maximal support according to the Shimodaira-Hasegawa test implemented in FastTree with 1,000 resamples. Branch lengths are in amino acid substitutions per site.

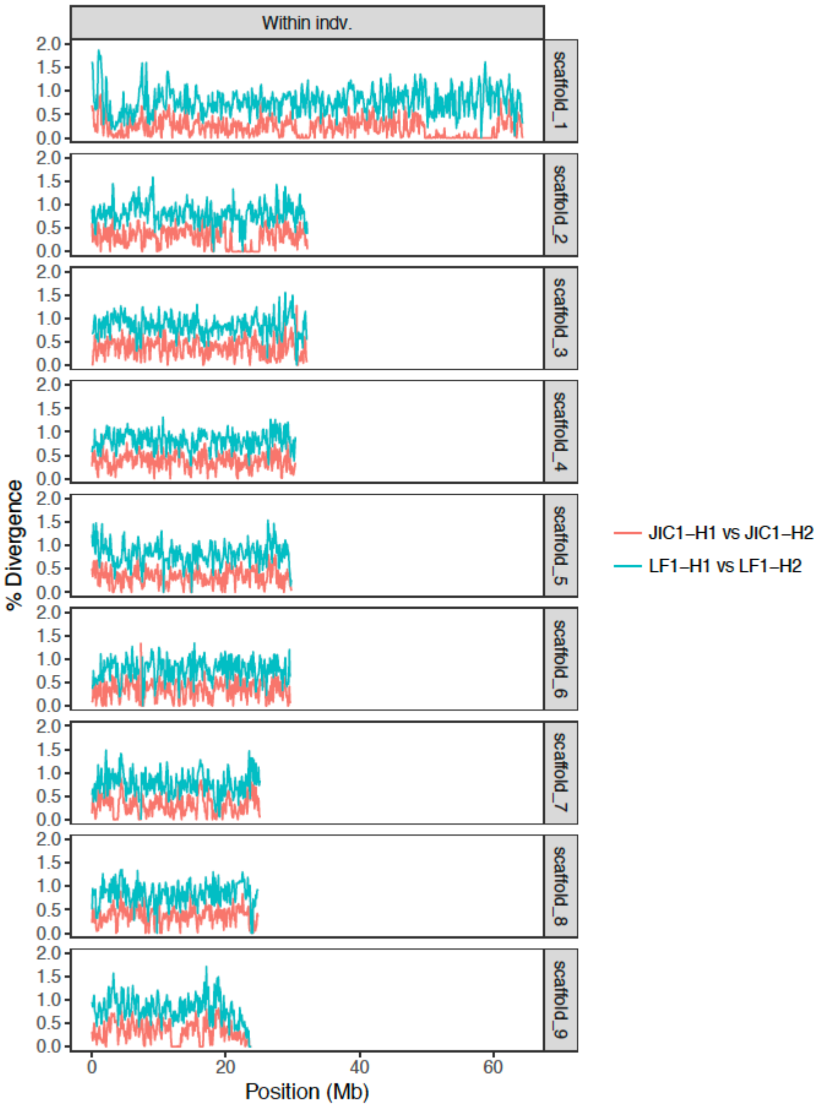


**Supplementary Figure 12:** JIC1 and LF1 within individual haplotype divergence (all chromosomes).

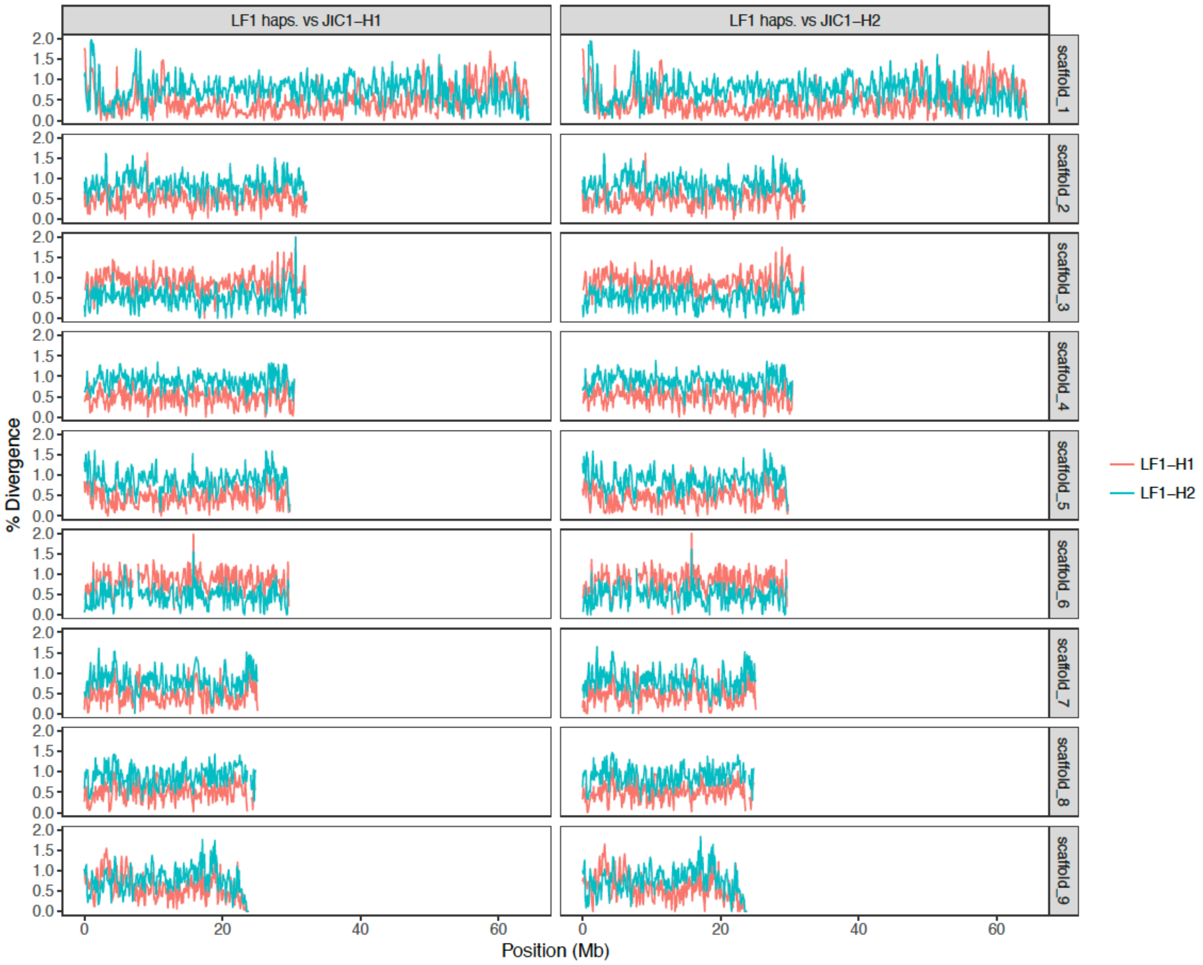


**Supplementary Figure 13:** JIC1 and LF1 between individual haplotype divergence (all chromosomes).

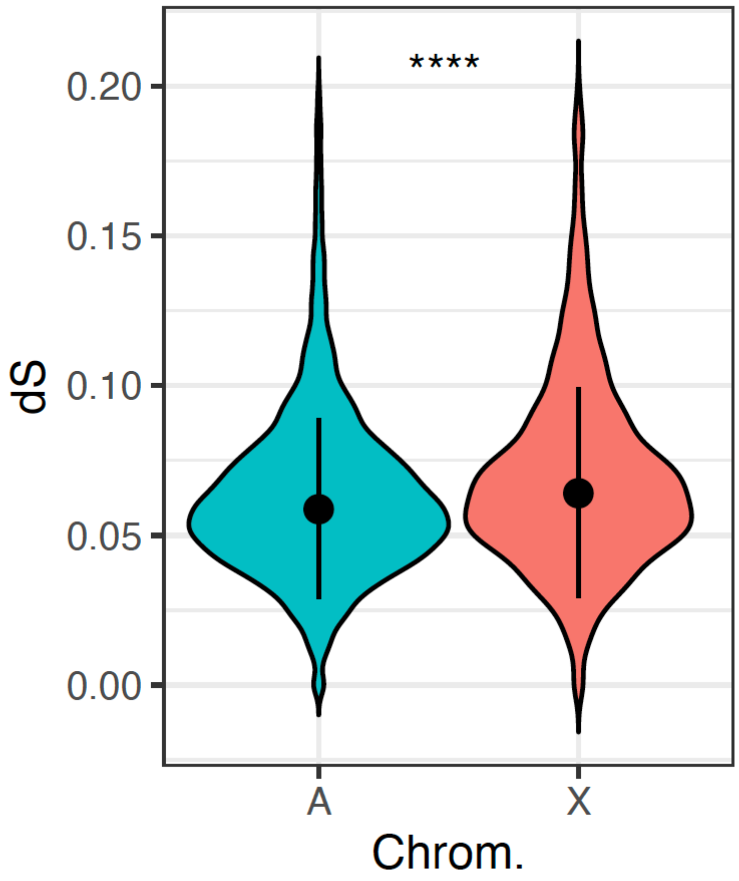


**Supplementary Figure 14:** *Violin plots* of synonymous site divergence (dS) between *S. miscanthi* and *M. dirhodum* one-to-one orthologs located on *S. miscanthi* autosomes (A; n = 8,462) and the *S. miscanthi* X chromosome (X; n = 1,871). Dot and whiskers show the median and interquartile range, respectively. X chromosome genes have significantly higher dS than autosomal genes (Wilcoxon rank sum test, $p = < 2.2x10^{-16}$, W = 8,879,400). Ortholog pairs with extreme dS values (dS >= 0.2) were omitted.

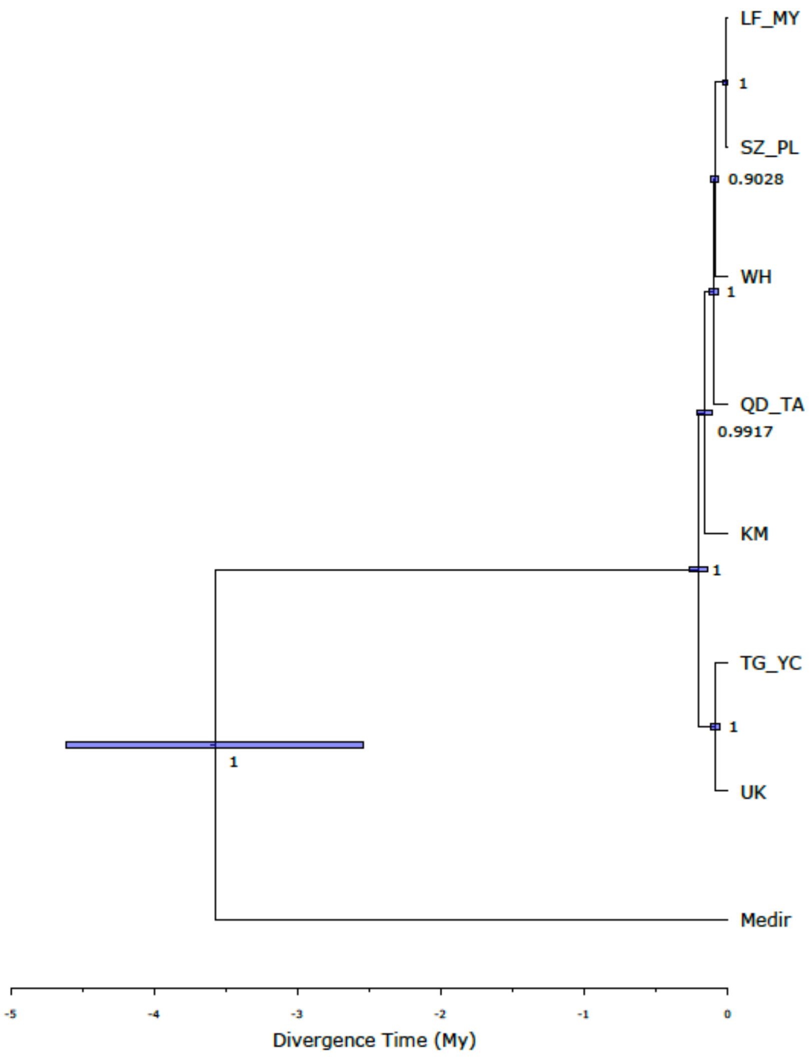


**Supplementary Figure 15:** SNAPP maximum-clade-credibility time calibrated phylogeny of *Sitobion* lineages rooted with *M. dirhodum* (Medir). Lineages are named according to **Figure 5** (*main text*). Bars at nodes indicate 95% highest posterior densities of date estimates. Numbers at nodes show posterior probabilities. My = millions of years ago.

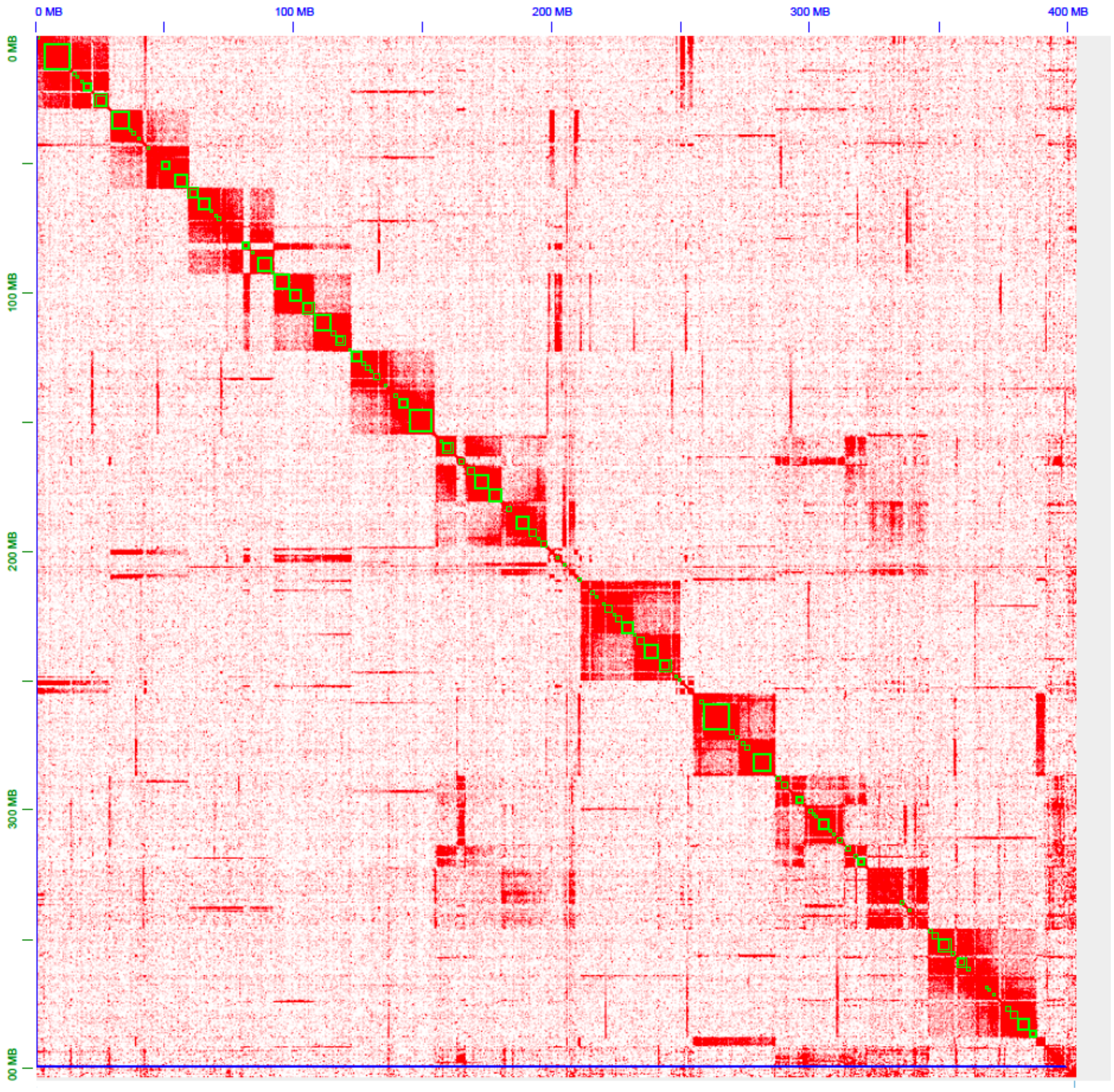


**Supplementary Figure 16:** Hi-C contact map for the draft *S. miscanthi* genome assembly after the initial round of scaffolding carried out by the 3D-DNA pipeline. The blue line shows the single super-scaffold assembled by 3D-DNA that is subsequently broken down into putative chromosome-scale scaffolds in later rounds of the pipeline. The green lines show assembly contigs. Due to poor scaffolding performance likely caused by low resolution of the Hi-C library, we manually edited the assembly in JBAT after this stage of the 3D-DNA pipeline to generate chromosome-scale scaffolds (see *main text* **Figure 1b**).

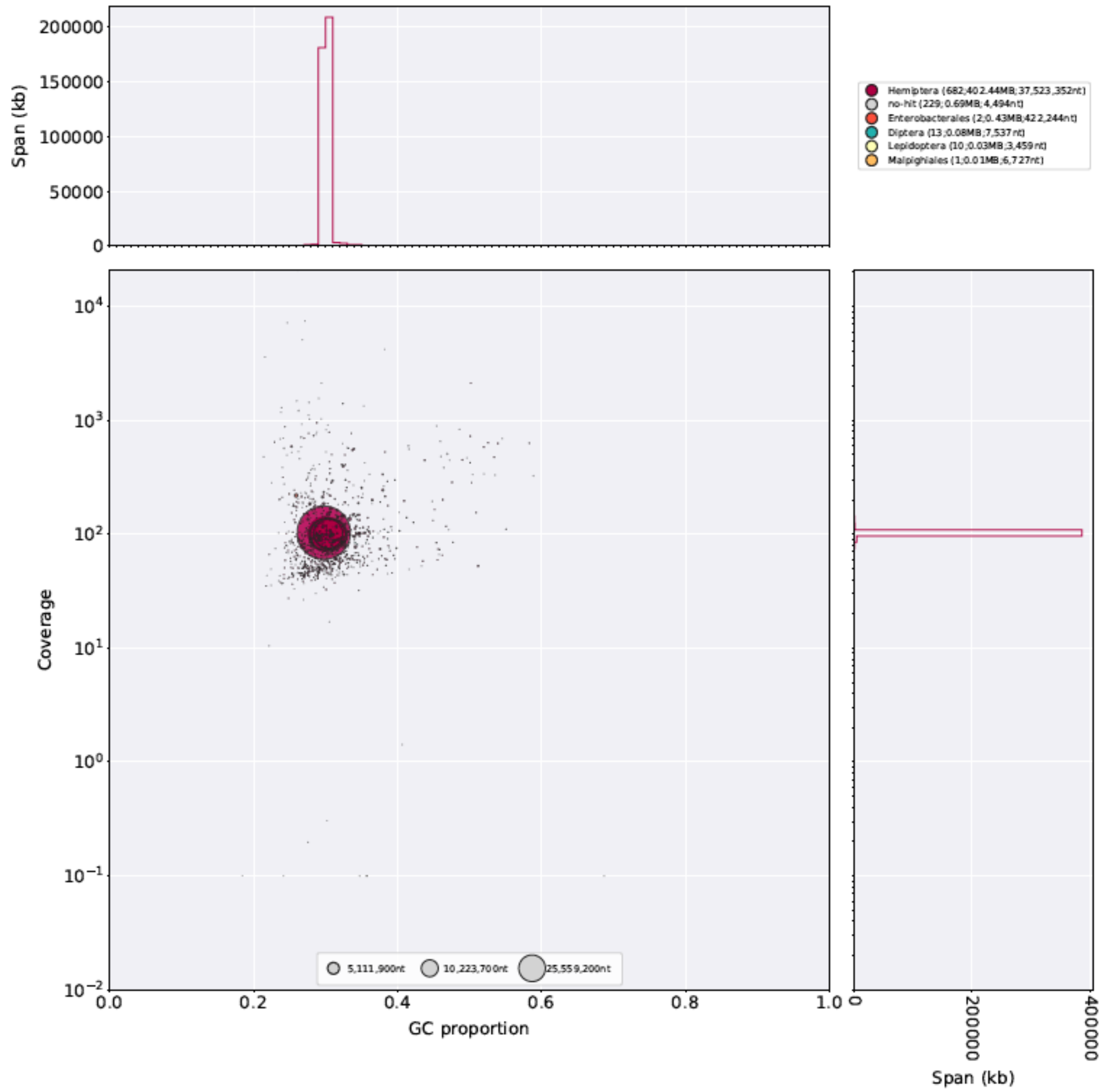


**Supplementary Figure 17:** Taxon-annotated GC content-coverage plot of the *Sitobion miscanthi* draft genome assembly after Hi-C scaffolding but before removal of symbionts and other contamination. Scaffold coverage (y-axis) is based on alignment of alignment Illumina paired-end reads from the *S. miscanthi* Langfang-1 colony (from Jiang *et al.* 2019). Taxonomy is annotated at the order level. See **Supplementary Figure 4** legend for detailed description of the plot.

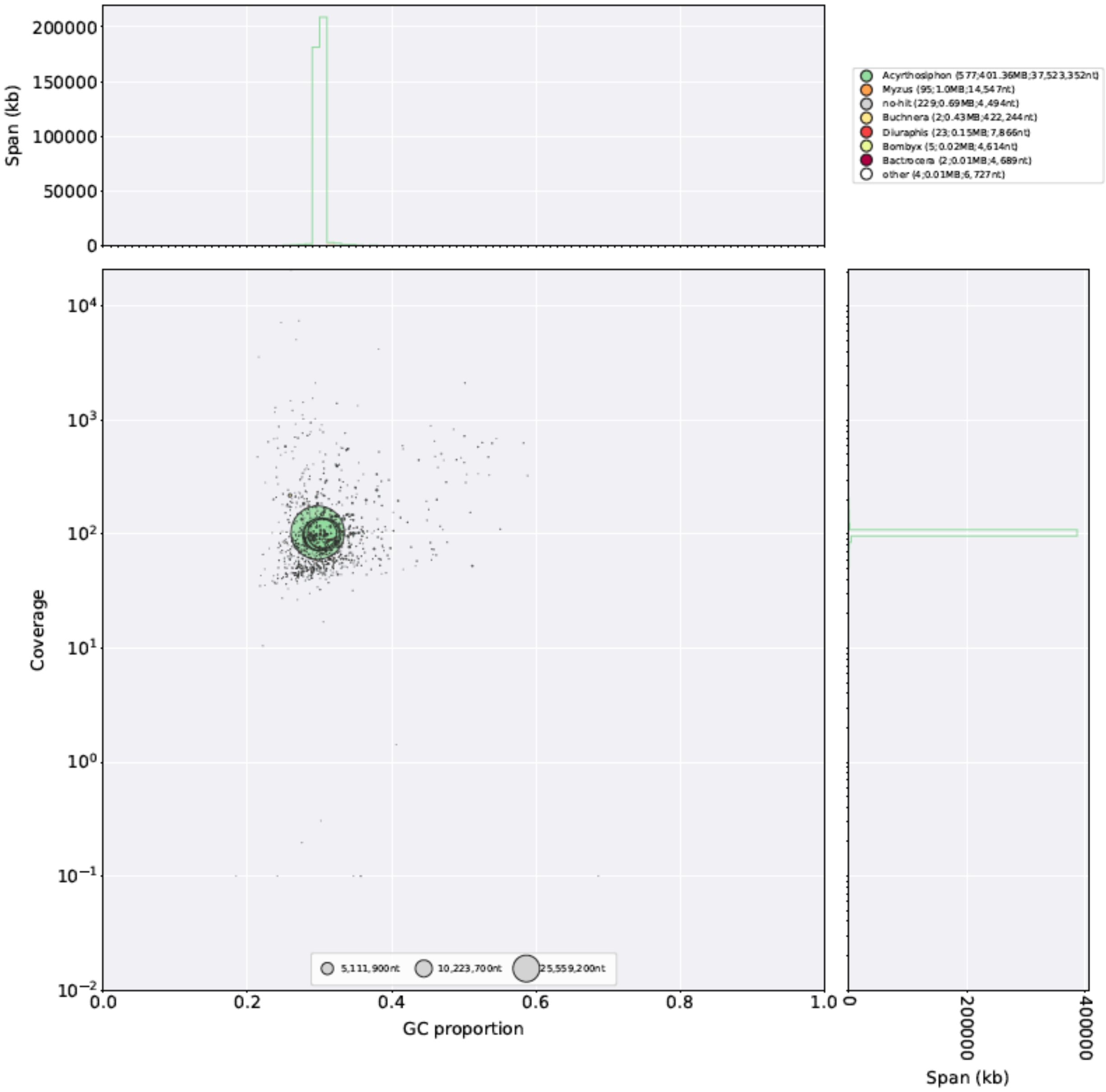


**Supplementary Figure 18:** As for **Supplementary Figure 17** but with taxonomy annotated at the genus level.

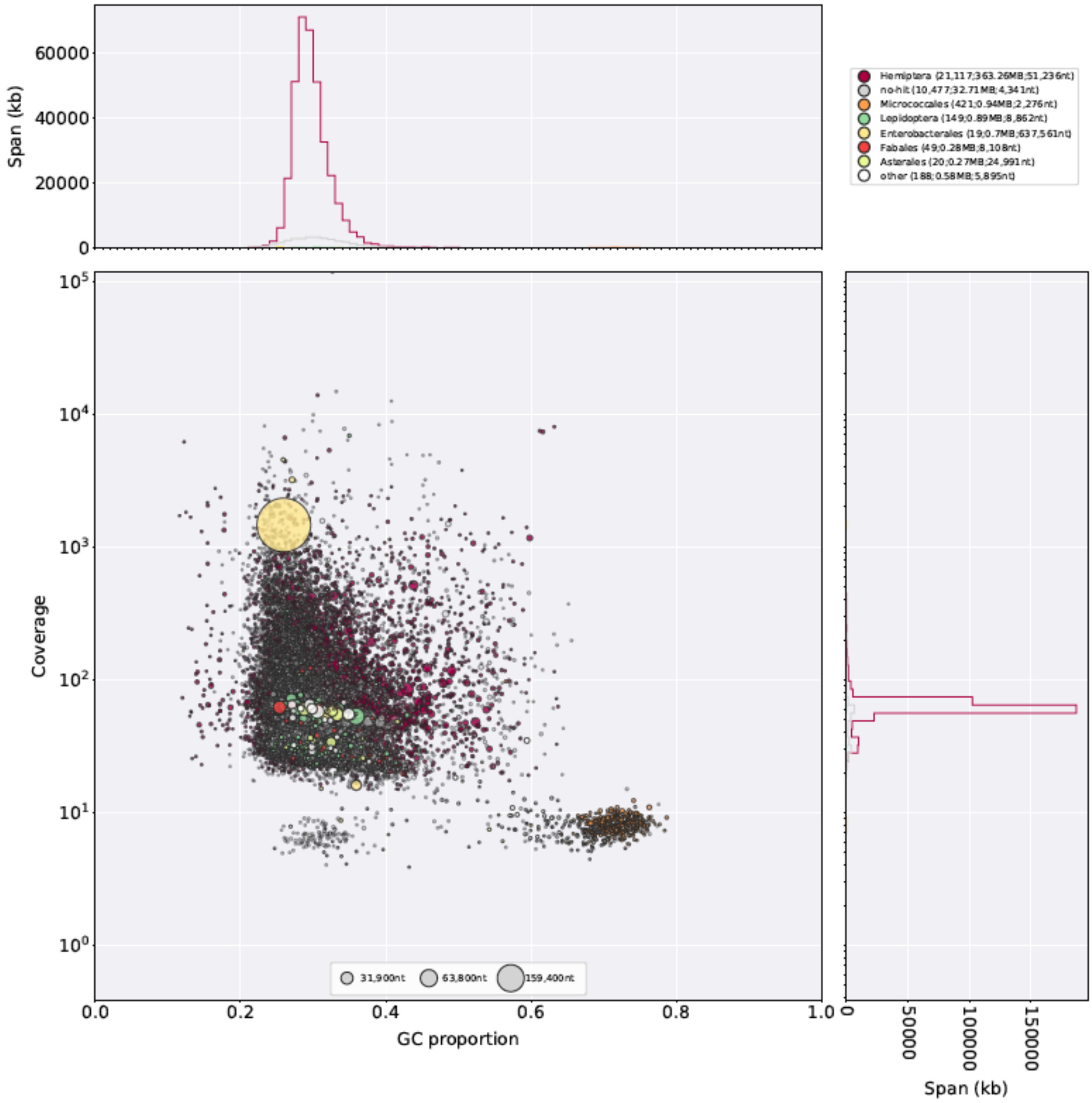


**Supplementary Figure 19:** Taxon-annotated GC content-coverage plot of the *S. avenae* Discovar *de novo* draft genome assembly after initial deduplication but before removal of symbionts and other contamination. Scaffold coverage (y-axis) is based on alignment of PCR-free Illumina paired-end reads used for the assembly. Taxonomy is annotated at the order level. See **Supplementary Figure 4** legend for detailed description of the plot.

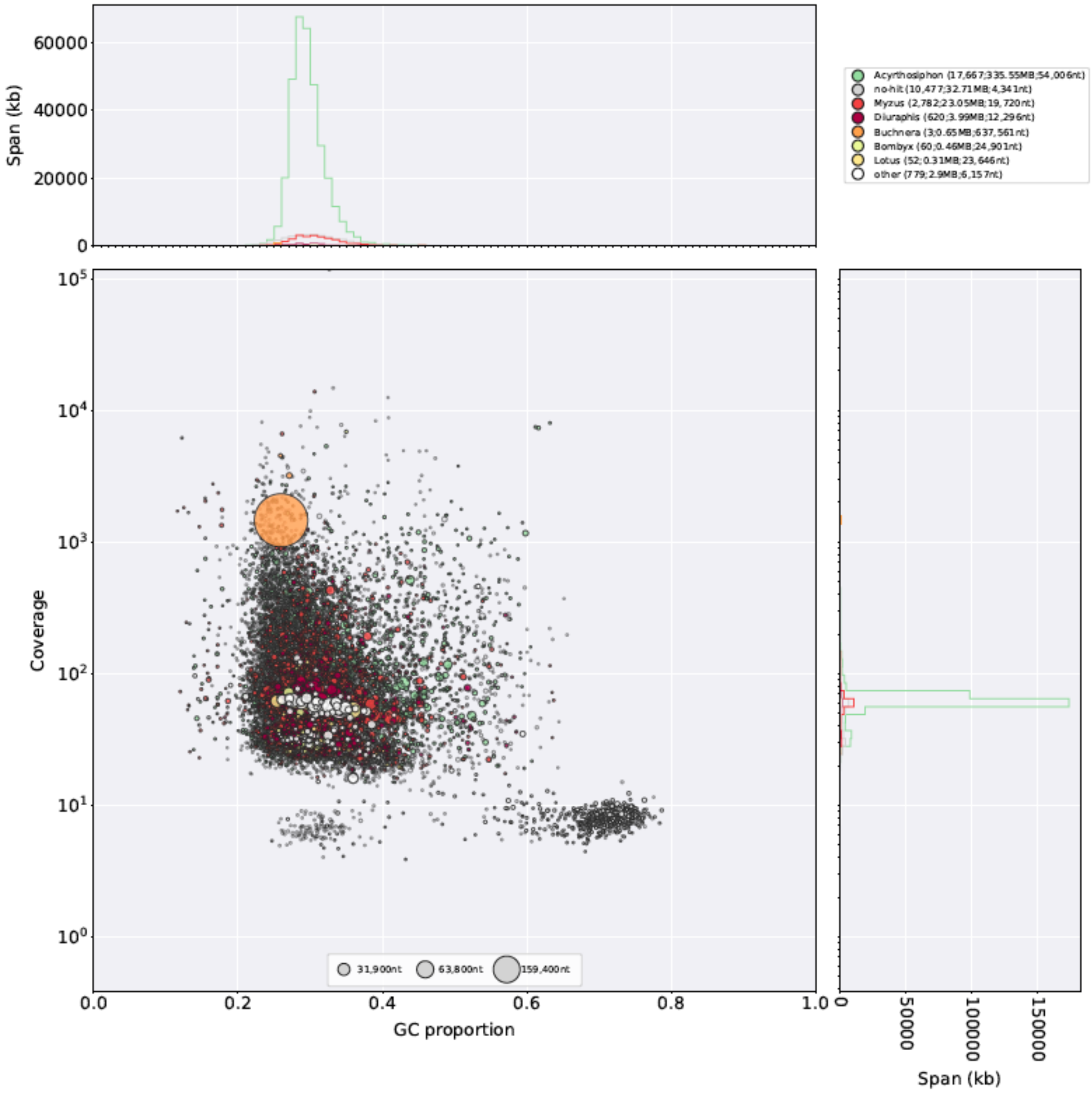


**Supplementary Figure 20:** As for **Supplementary Figure 18** but with taxonomy annotated at the genus level.

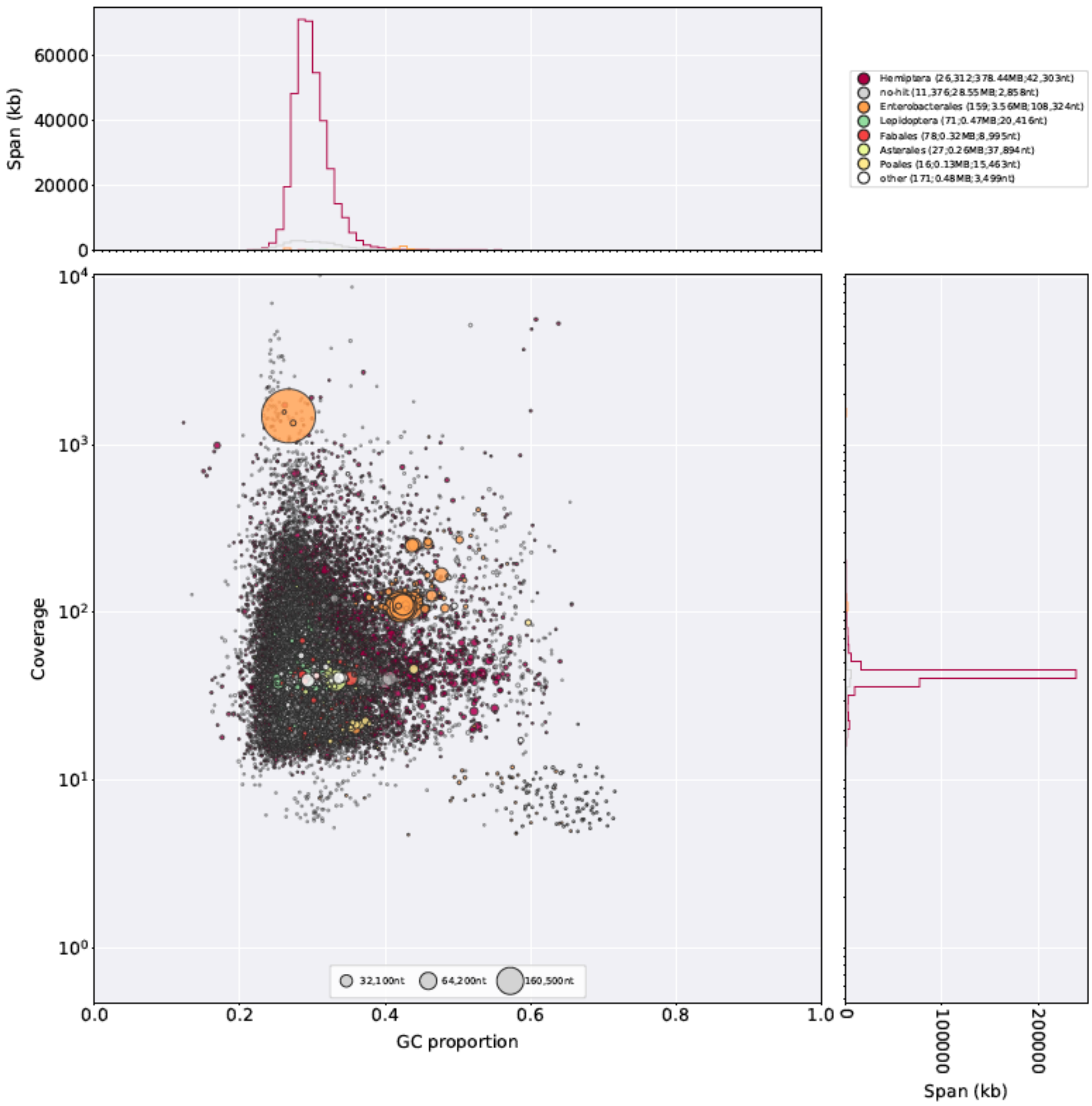


**Supplementary Figure 21:** Taxon-annotated GC content-coverage plot of the *M. dirhodum* Discovar *de novo* draft genome assembly after initial deduplication but before removal of symbionts and other contamination. Scaffold coverage (y-axis) is based on alignment of PCR-free Illumina paired-end reads used for the assembly. Taxonomy is annotated at the order level. See **Supplementary Figure 4** legend for detailed description of the plot.

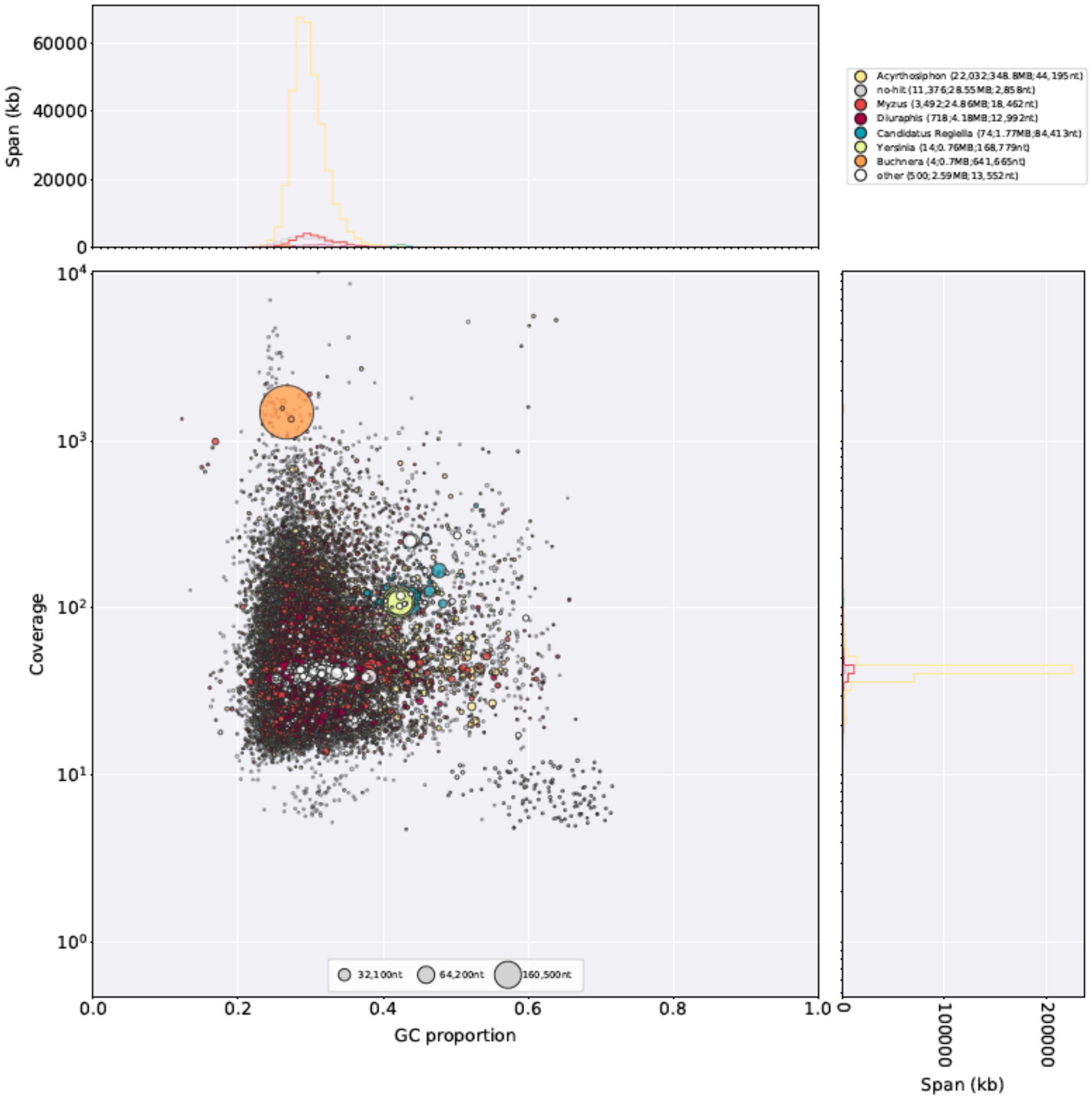


**Supplementary Figure 22:** As for **Supplementary Figure 21** but with taxonomy annotated at the genus level.

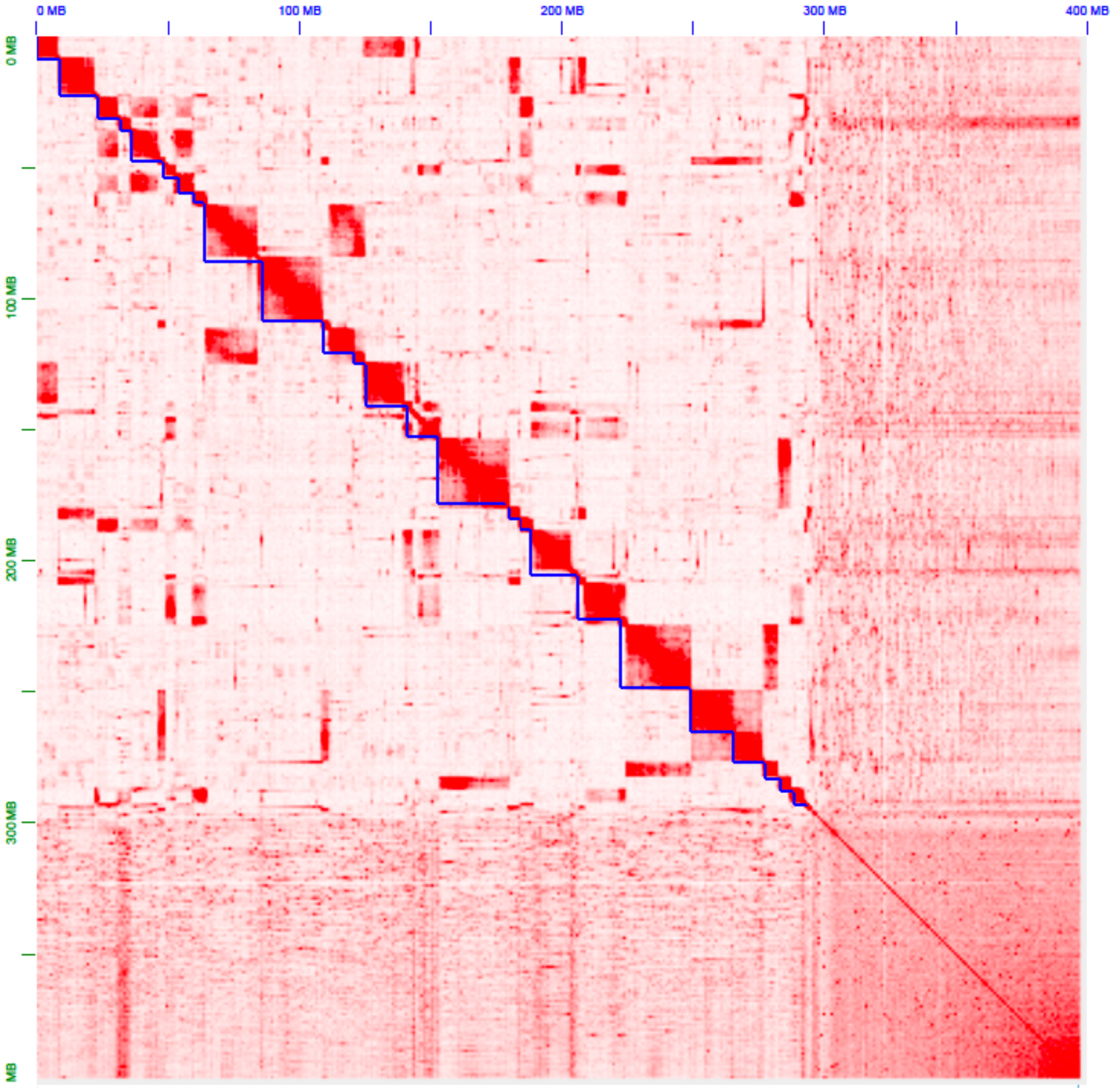


**Supplementary Figure 23:** Hi-C contact map for the 3D-DNA scaffolded draft *S. avenae* assembly before manual review with JBAT. Blue lines indicate super scaffolds.

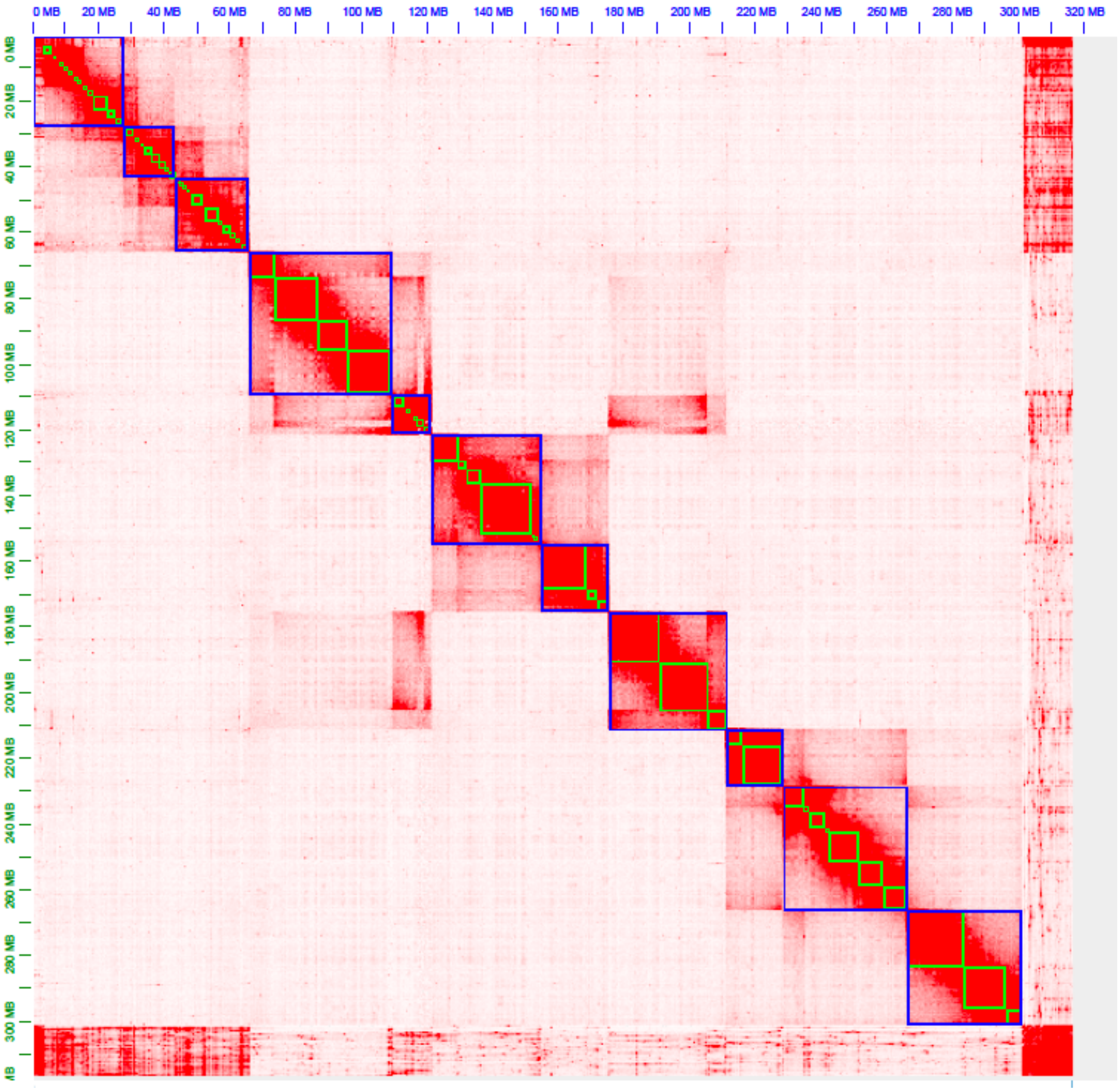


**Supplementary Figure 24:** Hi-C contact map for the 3D-DNA scaffolded draft *R. padi* assembly before manual review with JBAT. Blue lines indicate super scaffolds, green lines indicate scaffolds in the input draft assembly (Supernova + scaff10x + tigmint).

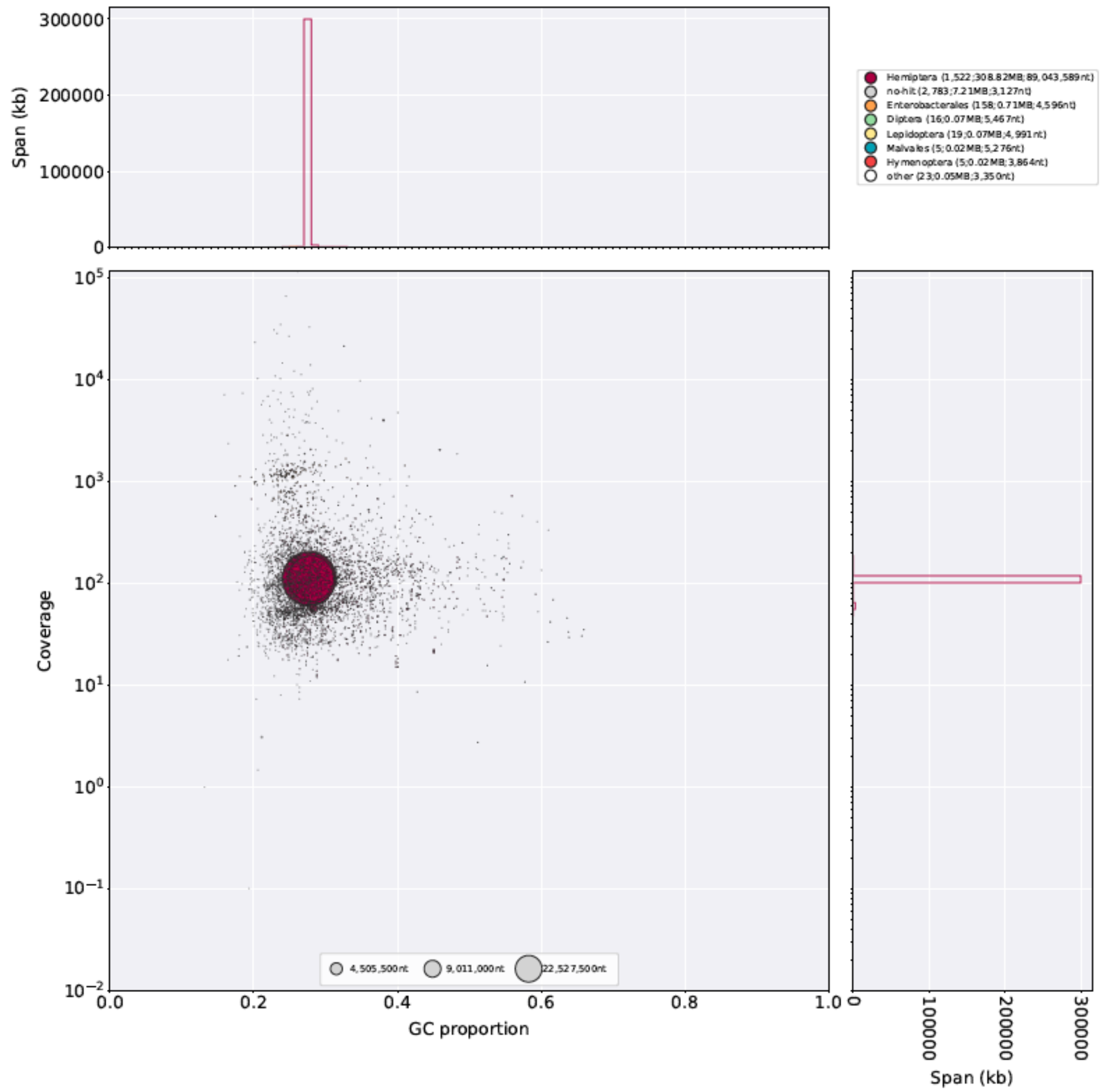


**Supplementary Figure 25:** Taxon-annotated GC content-coverage plot of the *R. padi* Discovar *de novo* draft genome assembly after after Hi-C scaffolding but before removal of symbionts and other contamination. Scaffold coverage (y-axis) is based on alignment of 10x genomics linked-reads used for the *de novo* assembly. Taxonomy is annotated at the order level. See **Supplementary Figure 4** legend for detailed description of the plot.

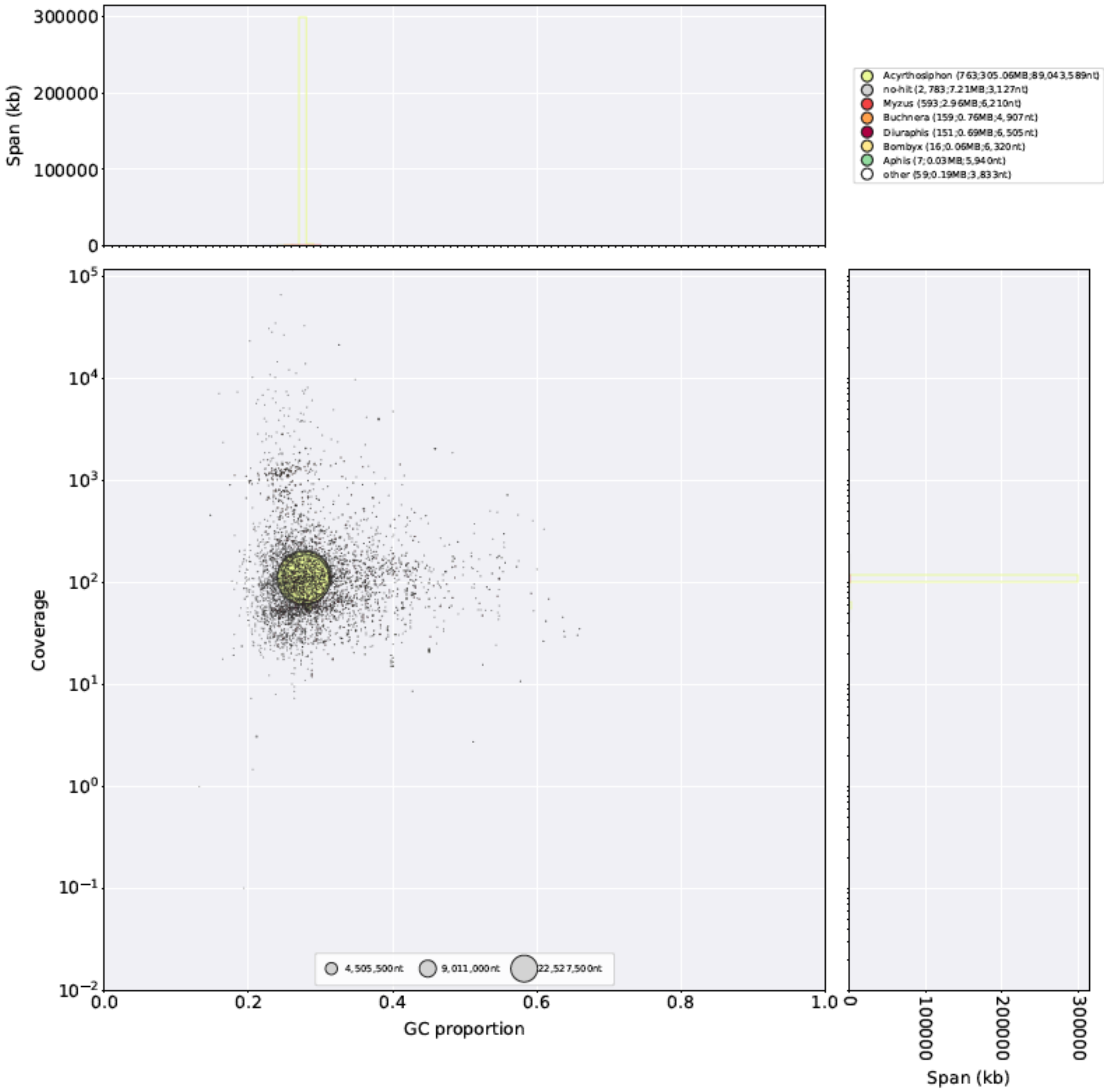


**Supplementary Figure 26:** As for **Supplementary Figure 25** but with taxonomy annotated at the genus level.
